# Small RNAs target native and cross-kingdom transcripts on both sides of the wheat stripe rust interaction

**DOI:** 10.1101/2022.06.16.496483

**Authors:** Nicholas A. Mueth, Scot H. Hulbert

## Abstract

The wheat stripe rust fungus (*Puccinia striiformis* f.sp *tritici*) poses a significant challenge to global wheat production. Plant defense induction against pathogens is partly modulated by small RNA (sRNA) molecules that downregulate complementary protein-coding transcripts. Additionally, the two-way RNA exchange between host and microbe involves cross-kingdom gene silencing that impacts pathogen virulence, yet few examples are known among rust fungi. The purpose of this study was to discover small RNA-target pairs on each side of this interaction. We performed sRNA sequencing and parallel analysis of RNA ends (PARE) in infected and uninfected wheat seedlings and combined these data with gene expression information. Wheat 24 nt heterochromatic siRNA (hc-siRNA) sequences were suppressed upon infection, while specific 35 nt tRNA and rRNA fragments were strongly induced. Target transcripts were identified by the observation of high mRNA slicing frequency at the precise position of sRNA binding sites. Wheat small RNAs showed evidence of cleaving several fungal transcripts including a ribosomal protein-coding gene and a glycosyl hydrolase effector gene. In *P. striiformis*, we confirmed and expanded previous findings that sRNAs feature microRNA-like sequences and siRNAs originating from long inverted repeats near protein-coding genes. Long inverted repeat regions produced sets of phased sRNAs at 21 nt intervals. Fungal sRNAs were identified that target native transcripts coding for transposons and protein kinases. Cross-kingdom gene targets of pathogen sRNAs included wheat nucleotide-binding domain leucine-rich repeat receptors (NLRs) and multiple families of defense-related transcription factors. The identified target genes shed light on an intricately co-evolved interaction and provide useful prospects for the development of pathogen control biotechnology.

## 1. Introduction

Wheat stripe rust is a destructive, globally distributed disease caused by the biotrophic fungus *Puccinia striiformis* f. sp. *tritici* [1, 2]. Wheat is cultivated on 220 million hectares worldwide [3]. Most of this area is continually or seasonally vulnerable to stripe rust; average annual losses exceed 5 million metric tons [4]. Stripe rust epidemics cause economically significant damage in the United States [5]. Timely fungicide application can mitigate losses, but has the disadvantages of added cost and environmental risk [6]. Deploying multiple resistance traits in wheat varieties to prevent, limit, or slow disease onset without using pesticides may be a more durable and sustainable strategy [7]. Moreover, basic research on trans-kingdom RNA interference (RNAi) has led to the development of host-induced gene silencing (HIGS) and exogenous spray-induced gene silencing (SIGS) as promising alternative strategies to control pests [8–10].

Plant-derived small RNAs play an important role in the defense against pathogens, especially secondary small interfering RNAs (siRNAs) that modulate the expression of immune receptor gene families [11]. Noncoding RNAs move within and between plants and fungi during host-microbe interactions [12]. The exchange of small RNAs between organisms is thought to be mediated by extracellular vesicles secreted by one interacting partner and taken up by the other [13]. Cross-kingdom silencing by small RNAs affects virulence, and can be manipulated to create more resistant or more susceptible phenotypes [14, 15]. Small RNAs 20-24 nt in length are present in most fungi and oomycetes, where they protect genome integrity, regulate development, and mediate interspecies interactions [16]. Small RNAs have been characterized from all three major *Puccinia* wheat rust fungi: *P. graminis, P. striiformis*, and *P. triticina* [17–19]. Recent work has advanced the genomic, transcriptomic, and epidemiological knowledge of *P. striiformis* [1, 20, 21].

Despite this progress, it remains difficult to confirm cases of cross-kingdom RNAi amid various native sRNA-associated processes in each organism. Computational approaches to sRNA-target prediction are prone to false positives, and often lack orthogonal evidence that the predicted target genes are actually expressed in the relevant tissues and conditions. To address this, small RNA-directed transcript slicing can be detected directly via parallel analysis of RNA ends (PARE). This technique generates sequencing tags from cleaved, degraded, or otherwise 5’ uncapped poly-A-containing transcripts, collectively known as the RNA “degradome.” Mapping PARE sequencing tags onto a transcript displays the frequency of fragments at each position. If a high-frequency peak is located at the 10th nucleotide of a small RNA binding site (the typical Argonaute-mediated slicing position), then this indicates a valid sRNA-target pair [22].

Gaining a complete picture of gene regulation in a host-microbe interaction necessarily involves examining the sRNA population, as well as both cleaved and intact mRNAs in both organisms. In this work, small RNA sequencing was combined with PARE and gene expression data to create an empirically-derived set of endogenous and cross-kingdom small RNA-target pairs in the stripe rust pathosystem.

## 2. Materials and Methods

### 2.1. Tissue collection

Wheat seeds of the variety “Avocet S” were germinated on moist filter paper for two days and then planted in 4×4-inch square soil pots at a density of 10-15 seedlings per pot. Seedlings were grown in a climate-controlled growth chamber (16 h light at 25 °C; 8 h dark at 15 °C). This cultivar is highly susceptible to *P. striiformis* strain PST-78. On the morning of the fourteenth day after planting, plants were hand-inoculated with PST-78 urediniospores diluted 1:10 with talcum powder (“Infected” treatment), or mock-inoculated with pure talcum powder (“Uninfected” treatment). Four individual pots composed independent replicates for the “Infected” treatment group, and four replicates for the “Uninfected” treatment. Pots were transferred to a dew chamber for 24 h in darkness at 10 °C and transferred to a climate-controlled greenhouse (~15 °C) for an additional seven days. Plants were then harvested just above soil level. Shoots within each pot were combined, immediately frozen in liquid nitrogen, and ground with a mortar and pestle. A few plants were left in each pot for observation for an additional two weeks to confirm heavy stripe rust sporulation in all inoculated reps and the lack of sporulation in uninoculated reps.

### 2.2. RNA extraction and RT-PCR

Total RNA was extracted from 1.0 g frozen ground tissue using TRIzol reagent (Thermo Fisher, USA). RNA quality (RNA integrity number > 6) was confirmed via Fragment Analyzer (Agilent, USA). RNA was treated with DNase I and reverse-transcribed using SuperScript III (Thermo Fisher, USA). RT-PCR was used to verify RNA quality and rule out cross-contamination between infected and uninfected samples. 30 cycles of endpoint PCR were performed using NEB Standard Taq (New England Biolabs, USA). PCR products (6 μL) were run on a 1% agarose gel with ethidium bromide for 20 minutes at 90 V and imaged under UV light. The wheat *GAPDH* transcript amplified from all samples, but fungal-specific Actin was exclusive to infected and not present in any uninfected replicate (**Supplementary File S1**). Oligonucleotide sequences can be found in **Supplementary File S2**.

### 2.3. Small RNA sequencing

The mirVana miRNA Isolation Kit (Thermo Fisher, USA) was used to extract small RNA (< 200 nt) from 200 mg frozen ground tissue. Small RNA libraries were prepared using the Illumina TruSeq sRNA system, and indexed barcodes specific to each rep were added for multiplex sequencing. Fragments were sequenced on four lanes of Illumina HiSeq at the Washington State University Genomics Core in Spokane, Washington.

Small RNA reads were mapped to the International Wheat Genome Sequencing Consortium *Triticum aestivum* RefSeq assembly v1.0 [23] using ShortStack version 3.8 [24]. Wheat plastid (NC_002762.1) and mitochondrial (NC_036024.1) genomes from NCBI GenBank were included. The following ShortStack options were used: --mismatches 0 --bowtie_m 250 --ranmax none --dicermax 40. Reads were also mapped to the Puccinia striiformis PST-78 draft genome (GCA_001191645) [25] using ShortStack. The following options were used: --mismatches 0 --mincov 50 --pad 200 --foldsize 400 --dicermax 40. ShortStack was used to discover phased siRNA loci using default settings. CLC Genomics Workbench 8.5 (QIAGEN, Netherlands) was used to process data. Wheat sRNA reads were identified as those mapping to the wheat genome, but not to the *Pst* genome. Fungal reads were identified as those mapping to the *Pst* genome, but not to the wheat genome. In uninfected libraries, a small fraction (~0.03%) of reads mapped exclusively to the fungal genome. Sequencing errors and index hopping can cause crosstalk between samples in multiplexed Illumina libraries [26, 27]. To prevent misidentification, in addition to mapping exclusively to the *Pst* genome, sequences attributed to the fungus were required either to be present exclusively in infected libraries, or else have an expression level >1,000-fold higher in infected vs. uninfected libraries. Sequences not meeting these criteria were excluded from downstream analysis.

RNA secondary structure prediction was calculated and visualized using CLC Genomics Workbench 8.5 and strucVis [28]. BLASTN was used to search for additional copies of *Pst*-MIRNA precursors in the genomes of *Pst* and other *Puccinia* spp. A positive match was defined as a hit with >80% sequence identity and >80% coverage of the originally-discovered MIRNA precursor query sequence, yielding an E-value < 1E-20, and in which the exact mature miRNA sequence was also found. The cutoff for identifying phased sRNA (phasiRNA) loci was a ShortStack cluster with phase score ≥ 50.

### 2.4. Parallel Analysis of RNA Ends

Total RNA was extracted with TRIzol from aliquots of the same frozen tissue used in RT-PCR and small RNA-seq. PARE libraries were constructed using the method of Zhai et al. [29] and sequenced on an Illumina HiSeq at Molecular Research LP (USA). PARE sequencing tags were assigned to each organism in a manner analogous to the small RNA reads. PARE reads that mapped to wheat cDNA sequences [24] but not to PST-78 cDNA [25] were considered wheat-derived. Fungal-derived PARE reads mapped perfectly to PST-78 cDNA, but not to wheat cDNA. PARE data were analyzed using CleaveLand 4 software on default settings using sRNA query sequences, PARE reads, and cDNA sequences as inputs [30]. The top 5,000 highest-expressed *Tae*-sRNA sequences 20-26 nt long, plus the set of all mature *T. aestivum* microRNA sequences from miRBase version 22 [31] were used as wheat query sequences. Fungal sRNA queries were the top 5,000 highest-expressed 20-26 nt *Pst*-sRNAs, plus the mature *Pst*-miR sequences discovered via ShortStack. The cutoff for positive CleaveLand results were as follows: the sRNA-target pair must be the highest peak for a given transcript (Category 0 or 1) in at least three out of four independent libraries. Measures were also taken to rule out the possibility that apparent cross-kingdom silencing is instead caused by an endogenous process. For *Pst*-sRNAs targeting wheat transcripts, the degradome density output files were checked to ensure that no significant peak (Category 0 or 1) was present at the same position in the uninfected treatment. Likewise, wheat vs. *Pst* PARE results were checked to rule out native *Pst* vs. *Pst* interactions.

Fungal target genes were annotated using the information at Ensembl Fungi [32] and UniProt [33]. Cleavage sites were annotated based on whether they appear in the predicted 5’UTR, coding sequence, or 3’UTR of the transcript. Protein sequences were searched for functional domains, including transmembrane and signal peptide domains, using HMMER searches against the Reference Protein database [34], with significant matches below an E-value cutoff of 0.01 (default setting). Protein homologs in other *Puccinia* spp. were identified via BLASTP searches of the EnsemblFungi database, with significant matches below an E-value cutoff of 1E-10.

### 2.5. RNA-seq data

RNA-seq reads were downloaded from a previous time course experiment of wheat seedlings infected with stripe rust at 0, 1, 2, 3, 5, and 7 days post infection [21]. The susceptible cultivar from that study was “Vuka”; the resistant cultivar was “Avocet” introgressed with the Yr5 stripe rust resistance gene. Illumina paired-end RNA-seq reads were mapped to the annotated genes in the wheat IWGSC RefSeq v1.0 genome or the *P. striiformis* PST-78 genome using CLC Genomics Workbench version 8.5 (Qiagen, Netherlands). Expression levels were calculated as the mean of three replicates at seven days post-infection; the units were reads per million mapped gene reads per kilobase (RPKM).

### 2.6. Statistical Analysis

Tests for differences between treatments (sRNA length, siRNA class, miRNA frequency, etc.) were as follows. Small differences in library size were corrected for by normalizing to the mean number of reads mapped to the wheat genome across all samples. The normalized count of small RNAs at each length was compared for infected vs. uninfected samples using Student’s t-test (n = 4 samples per treatment, α = 0.05). For analysis of known miRNAs, sRNA reads were mapped to *T. aestivum* precursor sequences from miRBase version 22 using CLC Genomics Workbench 8.5.

## 3. Results and Discussion

### 3.1. Wheat small RNAs

#### 3.1.1. Sequencing overview

High throughput Illumina sequencing generated over 650 million total sRNA reads (**Table 1**). 85% of total reads in the Uninfected treatment mapped with zero mismatches to the wheat genome, but not to the *P. striiformis* genome. The addition of fungal biomass/RNA caused the proportion of wheat-mapped reads to be lower in infected samples (66%). Approximately 38 million reads from infected samples (11% of total) mapped with zero mismatches to the fungal draft genome, but not to the wheat genome. The mapping and filtering procedure enabled partitioning of reads into sets of sequences originating from each organism.

**Table 1.** Small RNA sequencing and mapping summary. 18-40 nt sRNAs mapping to the wheat (*Triticum aestivum*) and *Pst* (*Puccinia striiformis*) genomes in uninfected and infected plants.

| Uninfected wheat | Reads | Unique sequences |
| --- | --- | --- |
| Total (n = 4) | 312,590,815 | 13,130,579 |
| Mapped exclusively to wheat genome | 265,393,738 | 7,723,099 |
| % | 85% | 59% |
| Infected wheat | Reads | Unique sequences |
| Total (n = 4) | 342,910,114 | 14,545,833 |
| Mapped exclusively to wheat genome | 225,379,229 | 6,163,742 |
| % | 66% | 42% |
| Mapped exclusively to *Pst* genome | 38,136,624 | 1,896,296 |
| % | 11% | 13% |

#### 3.1.2. Wheat sRNA size distribution

The size distributions of wheat small RNAs were plotted for total reads and unique (nonredundant) sequences. Infected wheat had significantly fewer 24 nt (13% of total) reads than uninfected wheat (16%) (**Figure 1A**). Fewer 24 nt sRNAs were also observed when counting only unique sequences (**Figure 1B**). The size distribution for unique sequences differs from total reads; 24 nt sequences are responsible for most of the diversity in the wheat sRNA repertoire, consistent with other angiosperms [35]. During stripe rust infection, many 24 nt wheat sRNAs disappeared or became dramatically under-expressed. This was previously observed in sRNA libraries made from wheat flag leaves of 6-week old wheat plants inoculated with stripe rust strain PST-100 [18]. In that study, the susceptible infected cultivar had fewer 24 nt reads (17% of total), compared with the susceptible uninfected control (20% of total).

**Figure 1.**
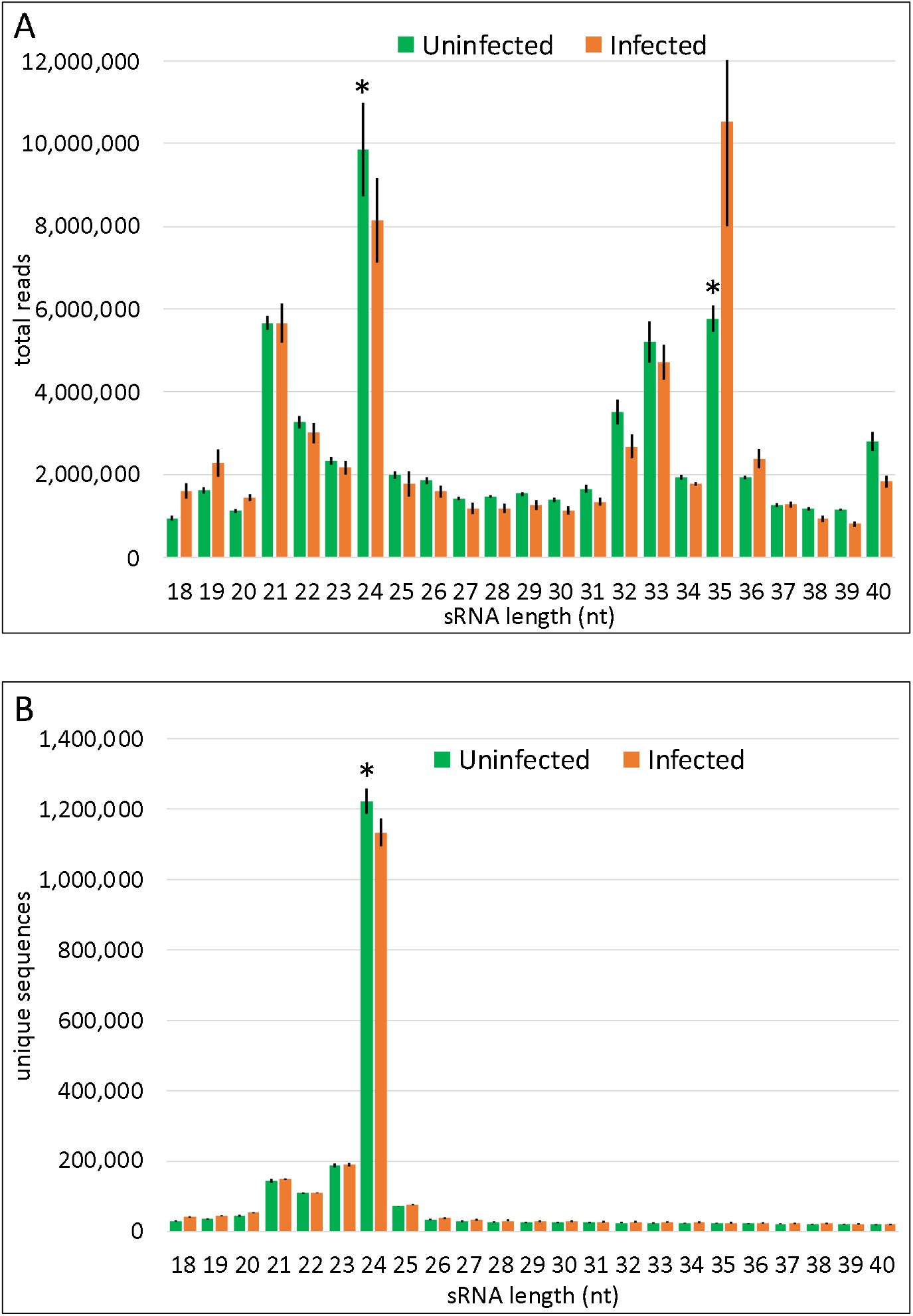
Wheat small RNA size distribution. Values are means for each treatment (n = 4). Error bars +/-1 SD. *Statistically significant (p < 0.05, Student’s t-test). (**a**) Total reads; (**b**) Unique sequences.

#### 3.1.3. Wheat 24 nt sRNA

24 nt sRNA reads were partitioned into various classes to determine if a specific class is responsible for this difference (**Table 2**). Mapping reads to wheat microRNA hairpin precursor sequences from miRBase version 22 revealed eight wheat *MIR* genes expressed in our samples that produce 24 nt mature miRNAs. However, there was no significant difference in 24 nt miRNA expression between treatments. The sRNAanno program was used to annotate the wheat genome with loci that generate various small interfering RNA (siRNA) classes, including heterochromatic siRNA (hc-siRNA) and phased siRNA (phasiRNA) [36]. The bulk of 24 nt siRNAs were hc-siRNAs; significantly lower 24 nt hc-siRNA expression in infected samples was mostly responsible for the overall difference between treatments (**Table 2**). In plants, 24 nt hc-siRNA sequences are associated with transcriptional silencing [37]. hc-siRNA precursors are transcribed by RNA Pol IV from repetitive intergenic regions such as transposable elements, made double stranded by RNA-dependent RNA polymerase RdR2, and processed into 23-24 nt siRNAs by dicer endonuclease DCL3. Mature hc-siRNAs then associate with AGO4 to guide sequence-specific deposition of repressive chromatin marks. Perhaps in the face of biotic stress, plant cellular resources normally used to control transposons are diverted to defense processes, resulting in a pause in hc-siRNA production. Alternatively, an effector from *P. striiformis* may be suppressing one or more siRNA biogenesis pathways that incidentally includes hc-siRNA. An effector from stem rust (*Puccinia graminis* f.sp. *tritici*) is a suppressor of silencing in heterologous expression systems [38]. There is also a pathogen-derived silencing suppressor in the *Arabidopsis-Phytophthora* pathosystem, although in that case 21 nt secondary siRNAs are suppressed [39]. 21 nt phasiRNAs are often associated with response to biotic stress, where they modulate expression of the *NB-LRR* resistance gene family [40, 41], yet this sRNA class was not significantly differentially expressed in our data. There were also fewer 24 nt phasiRNAs in infected wheat, but this made up only a small part of the overall difference in 24 nt sRNA abundance (**Table 2**). 24 nt phasiRNAs are mostly expressed in developing reproductive tissue and are present at low levels in seedlings [42]. It is unknown whether 24 nt phasiRNAs play some role in response to fungal infection, or if the difference is merely incidental to downregulation of 24 nt sRNA in general.

**Table 2.** Expression of wheat small RNA classes. Values are means for each treatment (n = 4). *Statistically significant (p < 0.05, Student’s t-test).

| Small RNA class | Uninfected mean | Infected mean | p-value |
| --- | --- | --- | --- |
| All 24 nt sRNA* | 9,843,900 | 8,153,513 | 0.0336 |
| 24 nt miRNA | 34,023 | 33,127 | 0.4252 |
| 24 nt hc-siRNA* | 6,463,276 | 5,144,865 | 0.0372 |
| 21 nt phasiRNA | 211,003 | 204,219 | 0.3539 |
| 24 nt phasiRNA* | 30,974 | 20,699 | 0.0007 |

#### 3.1.4. Wheat 35 nt sRNA

Significantly more 35 nt wheat reads were observed during infection (**Figure 1A**). Unlike the 24 nt reads, 35 nt reads made up a small number of unique sequences that was similar in both treatments (**Figure 1B**). Specific 35 nt sRNA sequences were induced upon infection (>1.5 fold higher in Infected vs. Uninfected; mean difference per sample >1,000 reads). The loci producing 35 nt sRNA matching these criteria overlapped with chloroplast-encoded tRNA, genome-encoded tRNA, and genome-encoded ribosomal RNA genes (**Figure 2A, Supplementary File S3**). tRNA-derived fragments were predominantly 5’ halves of mature tRNAs that were cleaved at the anticodon loop [43]. Ta-tRF1-5A is a 5’ tRNA half derived from the wheat chloroplast-encoded tRNA-Phe (**Figure 2B**). It was one of several tRFs that were differentially expressed in infected plants (**Figure 2C**) 3’ tRFs terminating in the TΨC loop were also observed. tRNAs carrying anticodons for specific amino acids were overrepresented: tRNA-Phe and tRNA-Leu made up over 90% of the total counts (**Supplementary File S3**). Accumulation of tRNA fragments is a broadly-conserved response to biotic and abiotic stress in yeast, plants and animals, with inhibitory effects on translation. Endoribonucleases that are usually inhibited by ligands and/or sequestered activate in the cytosol to produce tRNA-derived fragments that impair ribosomal function [44, 45]. 35 nt ribosomal RNA fragments were also overexpressed in infection (**Figure 2D**). The example rRNA small subunit in **Figure 2E**, ENSRNA050013838, is located on wheat Chromosome 1A. rRNA-derived fragments often corresponded to regions with a local stem-loop secondary structure, and are perhaps also the result of RNase activity [45].

**Figure 2.**
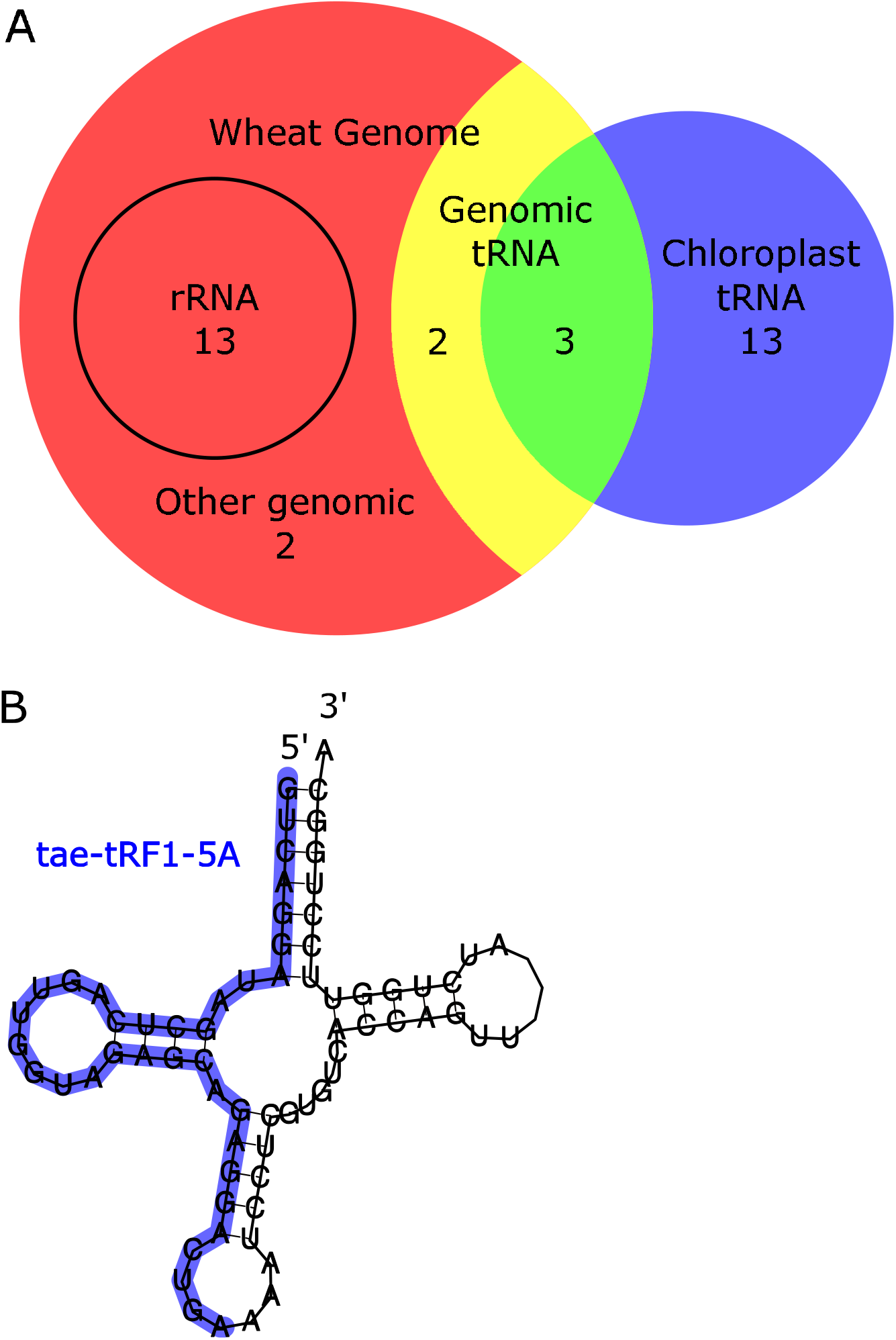
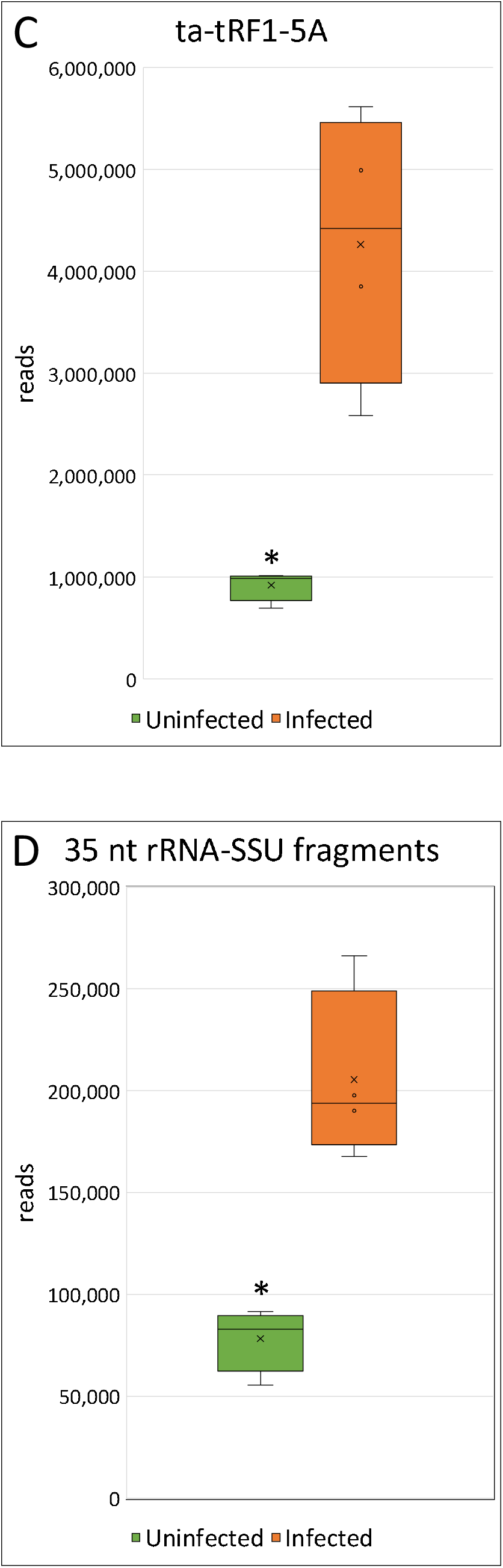
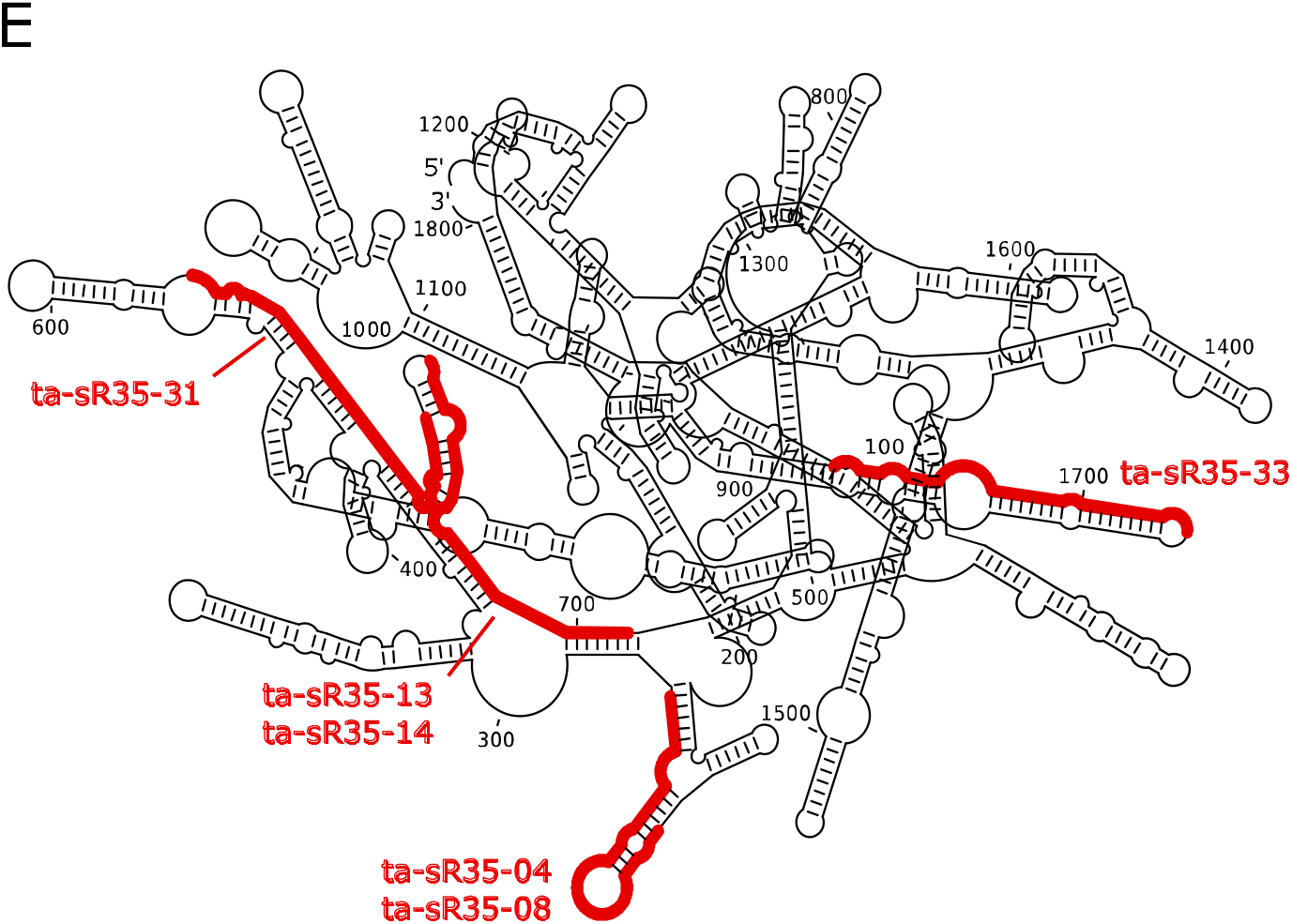
Highly-expressed 35 nt wheat RNAs induced during infection. (**a**) Fragments mapped to the wheat genome (large circle), or chloroplast genome (small circle). tRNA halves unique to the wheat genome (yellow), unique to chloroplast sequence (blue), or shared/ambiguous between plastid and wheat genomes (green); genomic rRNA fragments (red). (**b**) tRNA structure showing ta-tRF1-5A (blue), a 5’ tRNA half derived from the chloroplast-encoded tRNA-Phe. (**c**) Expression of ta-tRF1-5A in infected and uninfected samples. Values are means for each treatment (n = 4). *Statistically significant (p < 0.05, Student’s t-test). (**d**) Expression of 35 nt fragments derived from the wheat ribosomal RNA small subunit. (**e**) Infection-induced 35 nt fragments (red) overlaid on the rRNA-SSU structure.

### 3.2. P. striiformis small RNAs

#### 3.2.1 *Pst*-sRNA size distribution

The small RNA size distribution of *P. striiformis* is simpler than that of wheat. Sequenced reads consisted almost entirely of 19-23 nt sRNAs, with very few 24 nt or longer (**Figure 3A**). The size distribution is similar for both total reads and unique sequences (**Figure 3B**). 62% of 19-23 nt reads had uracil at the first position. This bias decreased for longer reads; only 34% of reads ≥ 24 nt began with 5’U.

**Figure 3.**
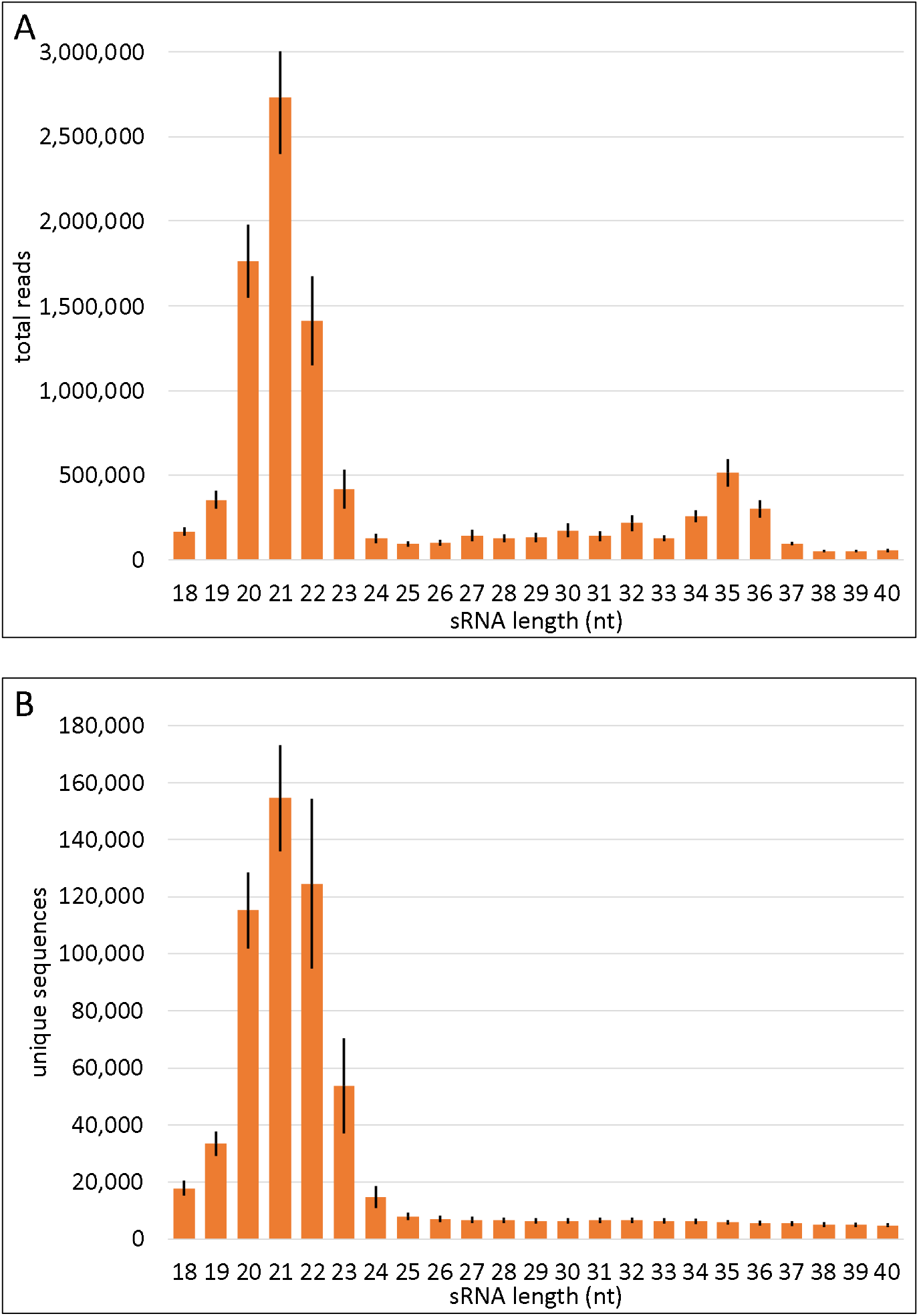
*P. striiformis* small RNA size distributions. Values are means for each treatment (n = 4). Error bars +/-1 SD. (**a**) Total reads; (**b**) Unique sequences.

#### 3.2.2. *Pst-*miRNA loci

We previously reported two microRNA-like loci in *P. striiformis* strain PST-100 [18]. These were also identified here in PST-78, indicating that they are conserved in at least two distinct strains. Increased sequencing depth enabled discovery of additional *Pst-MIRNA* loci (**Figure 4A**, **Supplementary File S4**). Precursors met all criteria for microRNA locus annotation, including presence of the miRNA/miRNA* duplex offset by 2 nt, limited mismatches/bulges between miRNA and miRNA*, and replication in multiple sequencing libraries [46]. All except *Pst*-MIR7 had more mature miRNA reads on the 5’ side of the hairpin than on the 3’ side. A majority of miRNA and miRNA* sequences began with 5’ uracil (**Supplementary File S4**). BLAST analysis of full-length MIRNA sequences found 1-5 copies of each in the PST-78 draft genome. At least one precursor homolog was also found in the draft genome for strain PST-130. *Pst*-MIR2 had sequence similarity to two loci in the *P. graminis* genome, but the region matching the mature miRNA sequence had several substitutions. No other MIRNA precursor had significant hits to any *Puccinia* species in the EnsemblFungi database (*P. coronata, P. graminis, P. sorghi*, or *P. triticina*). This suggests that *Pst-MIRs* are shared among intraspecific strains but are largely confined to stripe rust.

**Figure 4.**
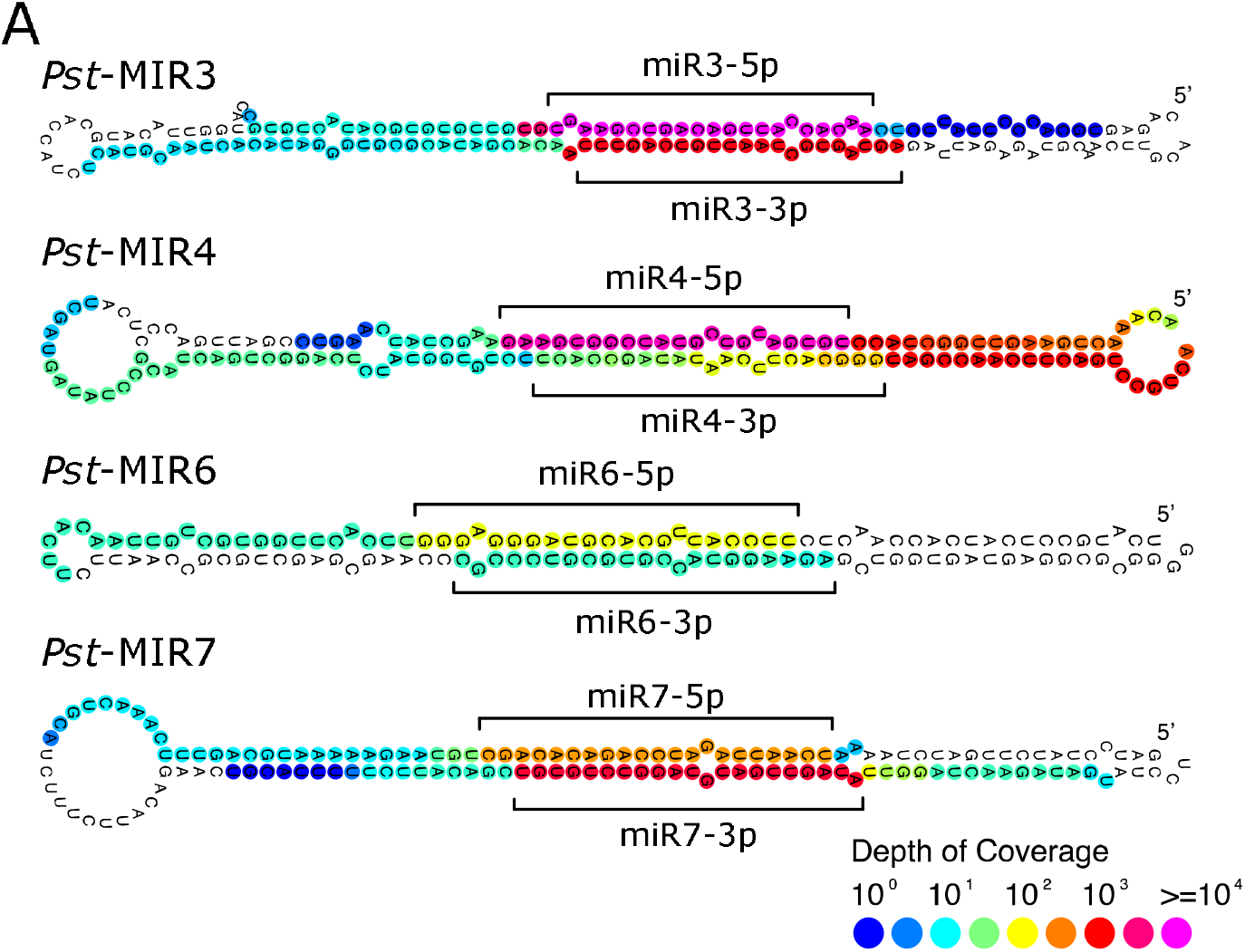
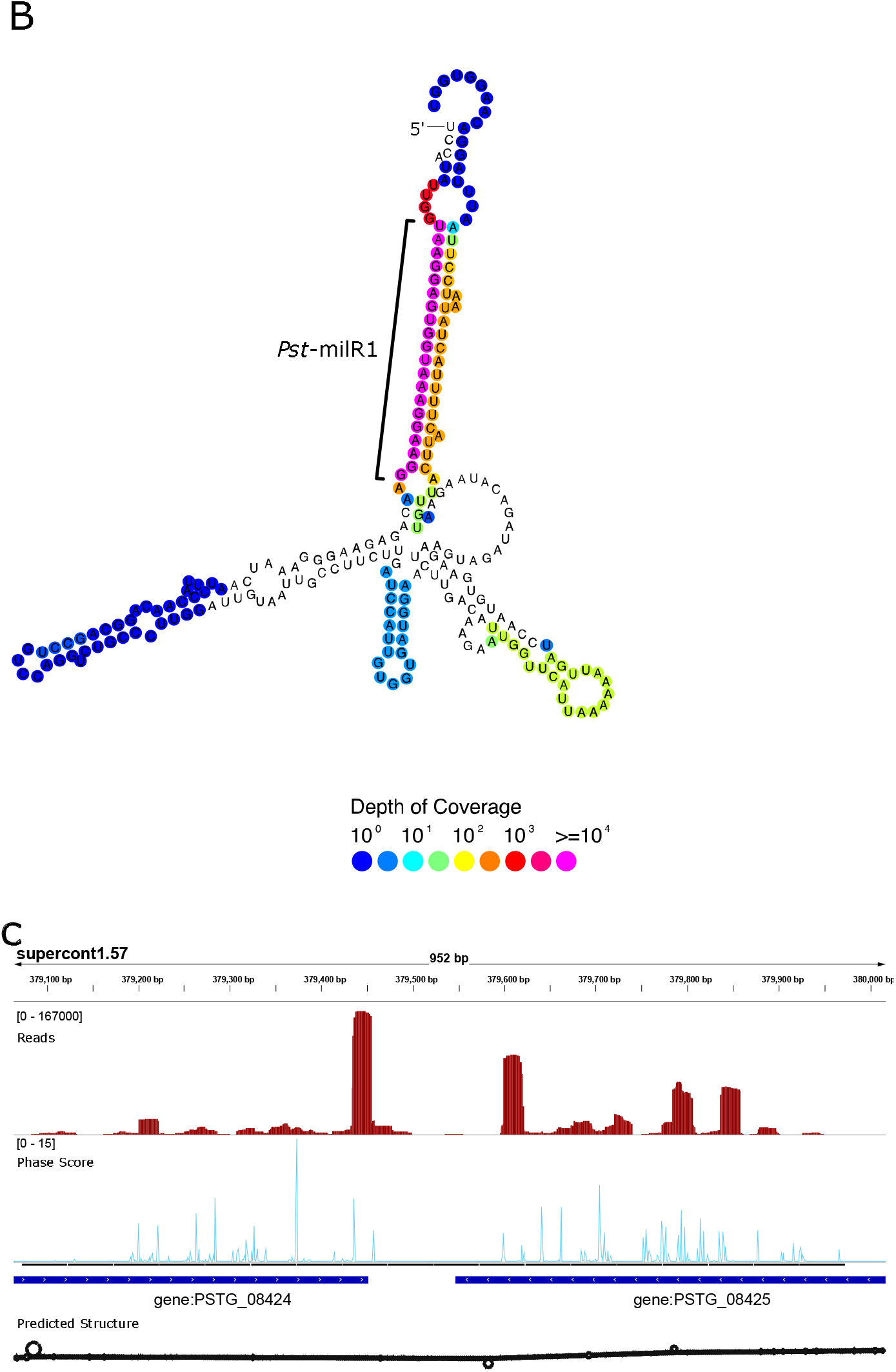
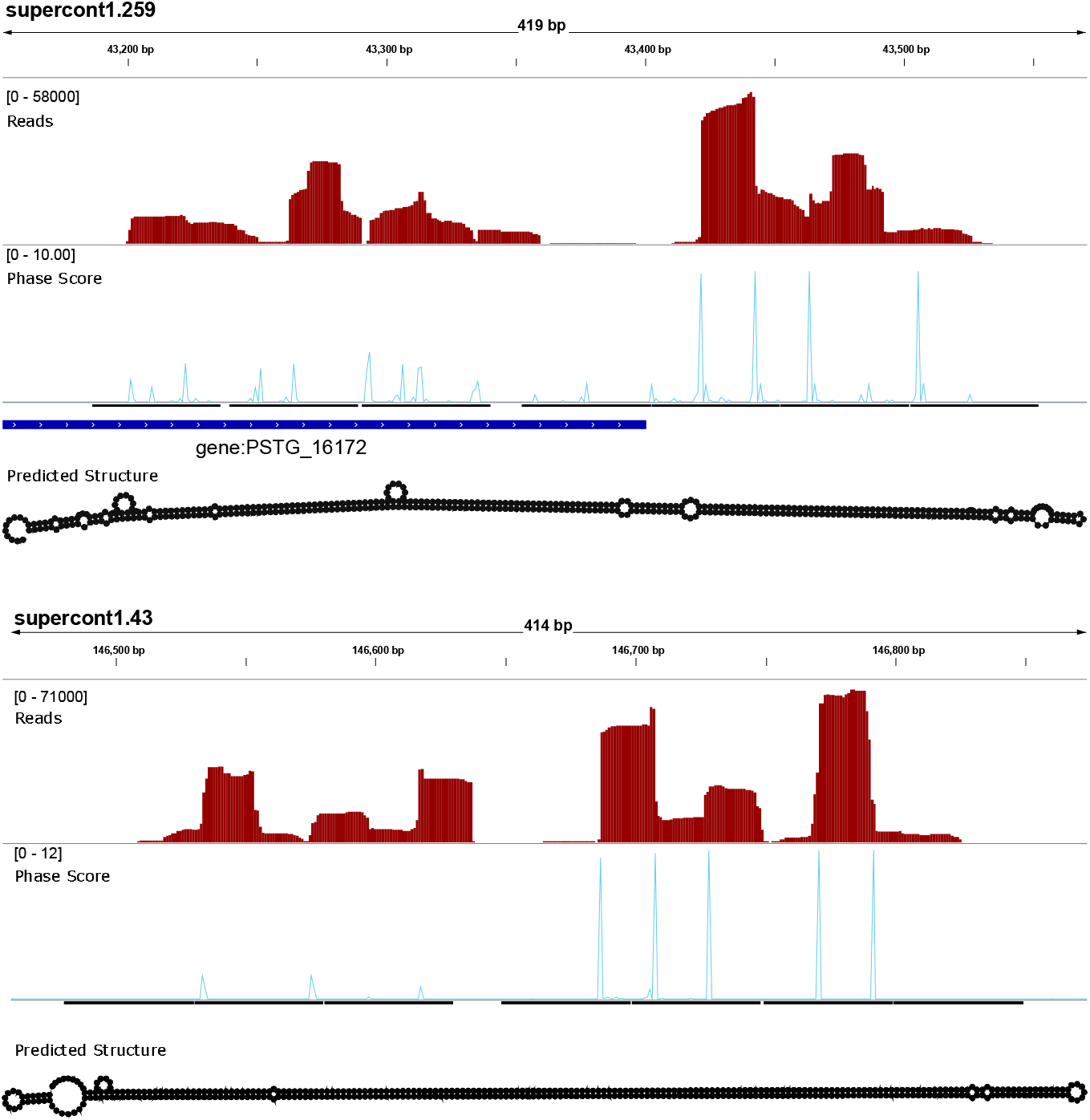
microRNA and small interfering RNA-producing loci in the *P. striiformis* genome. (**a**) Predicted *Pst*-MIRNA precursors. 5’ ends are all oriented top right. Read depth indicated by color. Mature miRNA-5p/miRNA-3p sequences indicated by brackets. (**b**) Predicted precursor of a locus producing *Pst*-milR1 described in [47]. (**c**) Long inverted repeat loci producing phased siRNA. Genome browser output showing read depth (red); phase score (light blue); gene annotations (dark blue); hypothetical RNA hairpin structure over the given interval (black). Top: the 3’ ends of a naturally antisense tail-to-tail gene pair produce siRNAs including *Pst*-milR1; middle: an inverted repeat flanking the 3’ end of a gene, produces 21 nt phasiRNAs; bottom: an additional long inverted repeat producing 21 nt phasiRNAs.

Wang and colleagues first described the stripe rust *Pst*-milR1, which was predicted to target a wheat transcript coding for a β-1,3 glucanase gene [47]. Reducing the abundance of *Pst*-milR1 using host-induced gene silencing (HIGS) correlated with a decrease in disease severity in wheat infected with *Pst* strain C YR31. The mature 20 nt *Pst*-milR1 (5’-UAAGGAGUGGUAAAGGAAGG-3’) was also abundant in our dataset, with over 18,000 reads in the total infected library. However, its precursor did not meet the criteria for annotation as a MIRNA locus due to a large asymmetrical loop within the precursor structure (**Figure 4B**). Also, the expected 20 nt miRNA* sequence with a 2 nt offset on the other side of the hairpin was not present in our dataset.

#### 3.2.3. *Pst*-sRNA from long inverted repeats

In addition to its previously reported precursor, mature *Pst-*milR1 maps to a separate locus in the PST-78 genome spanning the 3’ ends of two consecutive genes in a tail-to-tail arrangement (**Figure 4C**). If this inverted repeat region were transcribed in one piece, its RNA secondary structure would be a hairpin > 900 nt long. The locus produced a range of 20-24 nt long sRNA with reads mapping to both DNA strands. *Pst-*milR1 matches the protein-coding sequence of *PSTG_08424* and the 3’ UTR of *PSTG_08425*. Both genes code for uncharacterized hypothetical proteins with no recognizable domains or gene family annotations. Publicly-available RNA-seq data show no evidence for gene expression at 7 days post infection of *PSTG_08424* (0.0 RPKM), and low-level expression of *PSTG_08425* (2.14 RPKM) [21]. Despite the 3’ ends of the two genes being almost identical, the predicted peptide sequences are dissimilar (**Supplementary File S5**).

Many additional sRNA-producing loci in the *P. striiformis* genome take the form of long inverted repeats spanning the 5’ or 3’ ends of one or more predicted genes. These loci range in length from a few hundred to a few thousand basepairs. Small RNA reads in these regions have two possible mapping locations, one on each side of the inverted repeat. Thus, it is difficult to determine the unambiguous abundance pattern of mapped reads across the locus. It is also unclear whether the true precursor sequence is transcribed as one long RNA hairpin, or results from a heteroduplex of two distinct transcripts from opposite DNA strands. Fifteen sRNA-producing long inverted repeat loci throughout the genome show an additional feature – evidence of “phased” 21 nt siRNA production (**Figure 4C**, **Supplementary File S6**) [48]. phasiRNA was also reported in the white mold fungus *Sclerotinia sclerotiorum* [49].

### 3.4. Parallel Analysis of RNA Ends

#### 3.4.1. PARE overview

Sequencing of PARE libraries yielded 127 million total reads (**Table 3**). 94% of reads from uninfected replicates mapped exclusively to wheat cDNAs (including 5’ and 3’UTRs), compared with 89% from infected replicates. The sequencing depth was sufficient to map reads to most wheat transcript entries in the EnsemblPlants database. Out of 133,863 unique sequences, 75% were represented by at least one mapped read. In all samples, a significant fraction of reads mapped to a single transcript: the chloroplast-encoded *rbcL* gene coding for the large subunit of Rubisco. In infected samples, 2% of total reads and 19% of unique sequences mapped exclusively to *P. striiformis* PST-78 cDNAs. Out of 20,688 stripe rust cDNAs, one or more PARE reads mapped to 58% of all transcripts.

**Table 3.** PARE sequencing and mapping summary. Sequencing tags mapping to the wheat (*Triticum aestivum*) and *Pst* (*Puccinia striiformis*) transcriptomes in uninfected and infected plants.

| Uninfected wheat | Reads | Unique sequences |
| --- | --- | --- |
| **Total (n = 4)** | 53,223,313 | 5,492,578 |
| **Mapped exclusively to wheat cDNA** | 49,833,918 | 4,108,836 |
| **%** | 94% | 75% |
| Infected wheat | Reads | Unique sequences |
| **Total (n = 4)** | 73,985,169 | 6,843,917 |
| **Mapped exclusively to wheat cDNA** | 65,970,622 | 3,710,876 |
| **%** | 89% | 54% |
| **Mapped exclusively to *Pst* cDNA** | 1,766,732 | 1,279,020 |
| **%** | 2% | 19% |

#### 3.4.2. Wheat miRNAs target wheat transcripts

PARE data were validated by identifying known miRNA-target pairs in wheat such as the conserved plant miRNAs miR160, miR164, and miR398, which cleave their known targets. (**Table 4**, **Supplementary File S7**). In *Arabidopsis*, the miR160 family targets multiple *Auxin Response Factor* genes, influencing many processes controlled by auxin [50] (**Figure 5A**). A wheat homolog of the miR398 family targets a Cu/Zn superoxide dismutase gene involved in response to reactive oxygen species [51] (**Figure 5B**). Wheat-specific miRNAs were also identified, such as tae-miR9676-5p, which targets a transcript coding for a protein with a predicted signal peptide and abhydrolase domain [52]. Overall, 77% of cut sites were found in the coding sequences of target transcripts, compared with 23% in the 5’UTR or 3’UTR (**Figure 6A**). Using publicly available RNA-seq data [21], the expression of all target genes was corroborated (**Supplementary File S8**). We conclude that the dataset is sufficient to capture native regulation of plant gene expression.

**Table 4.**
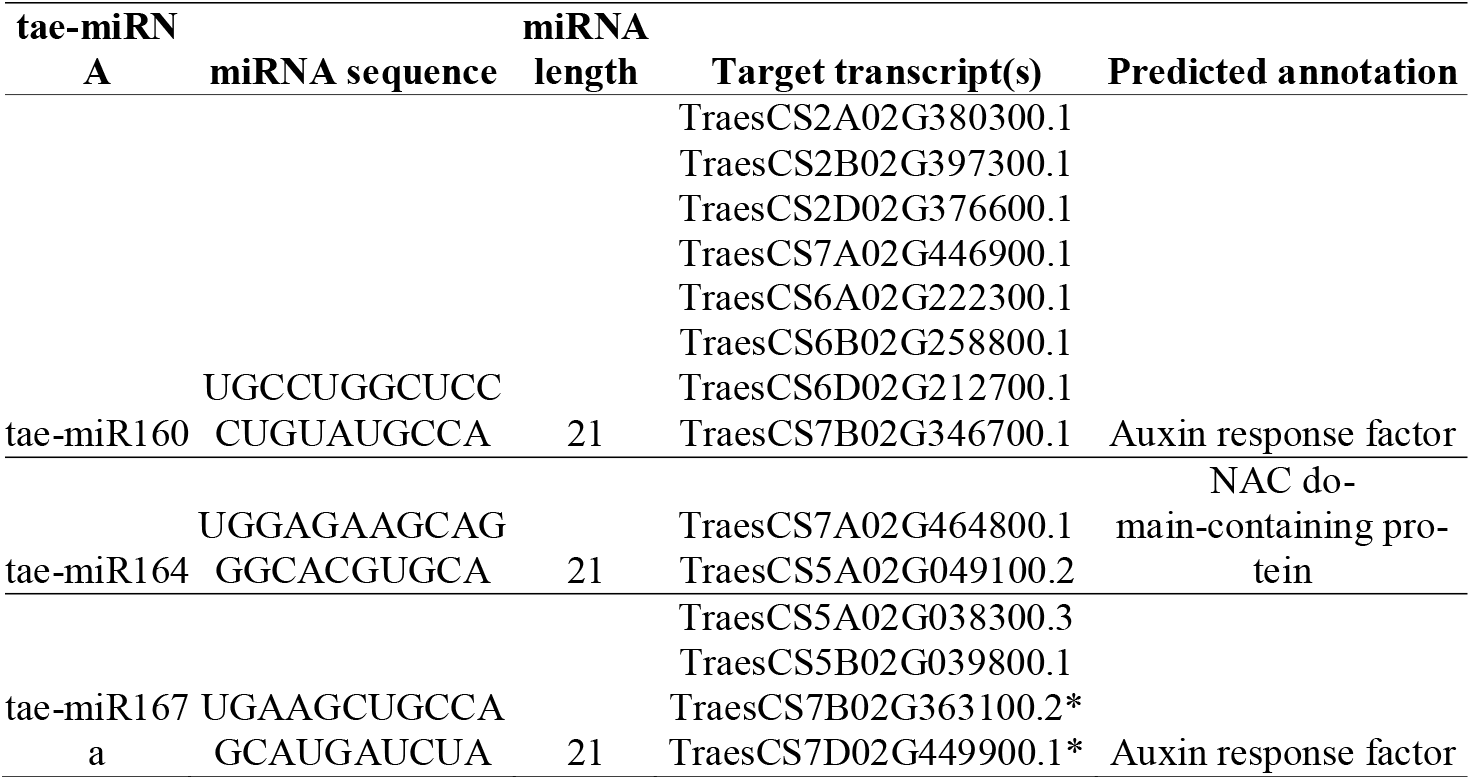
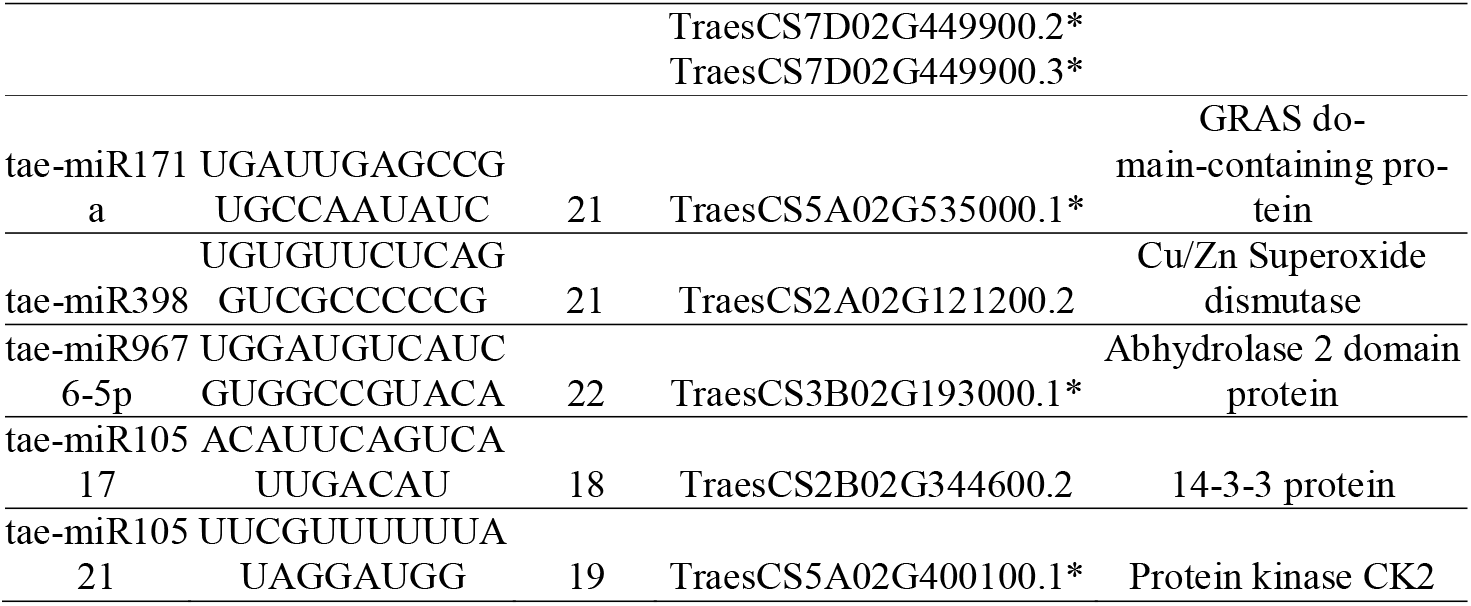
PARE: wheat sRNA vs. wheat transcripts. *Predicted protein contains a signal peptide or transmembrane topology.

**Figure 5.**
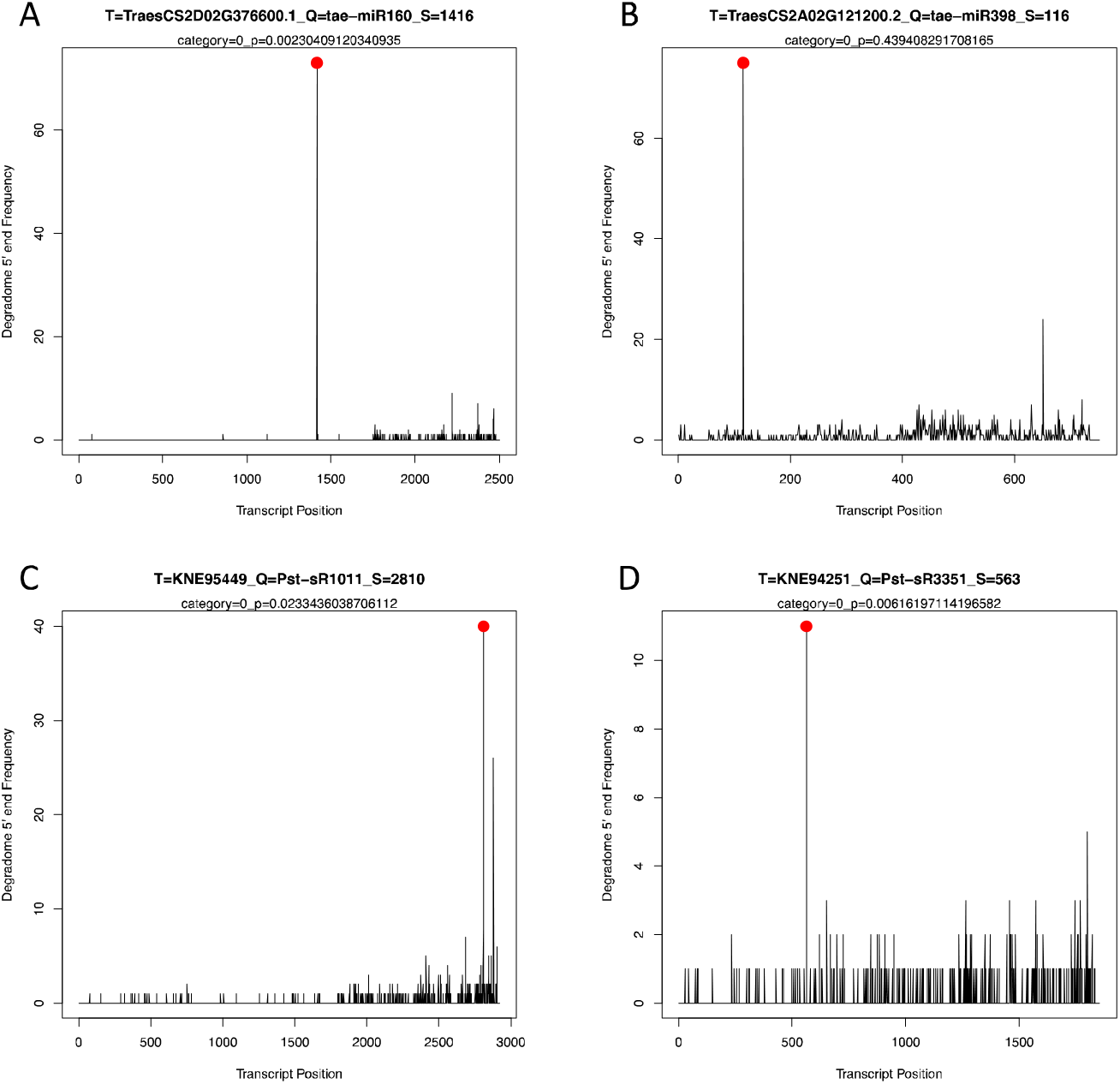
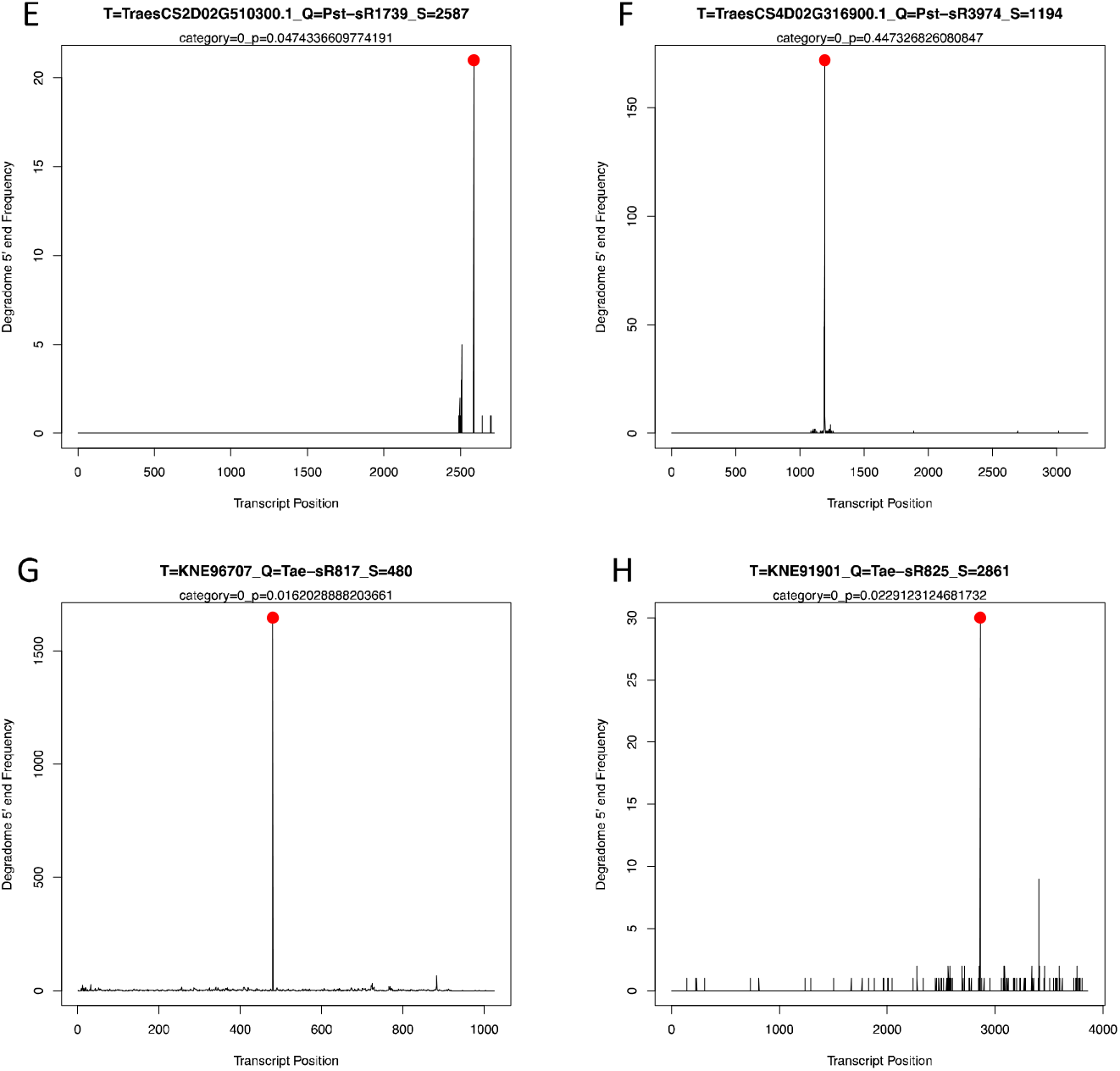
Target plots for select sRNA-target pairs. Peaks show the frequency of PARE sequencing tags at each transcript position. Red ball indicates the position of an sRNA-mediated cleavage site. Plots generated using pooled samples from infected plants (n = 4). (**a**) Wheat miR160 targets a predicted wheat auxin response factor transcript. (**b**) Wheat miR398 targets a predicted wheat Cu/Zn superoxide dismutase transcript. (**c**) *Pst*-sR1011 targets a predicted *Pst* transposase transcript. (**d**) *Pst*-sR3351 targets a predicted *Pst* cellulase transcript. (**e**) *Pst*-sR1739 targets a predicted wheat *NB-LRR* transcript (**f**) *Pst-*sR3974 targets a predicted wheat *bZIP* transcription factor transcript (**g**) Wheat tae-sR817 targets a predicted *Pst* ribosomal protein transcript. (**h**) Wheat tae-sR825 targets a predicted *Pst* TIMELESS domain-containing protein transcript.

**Figure 6.**
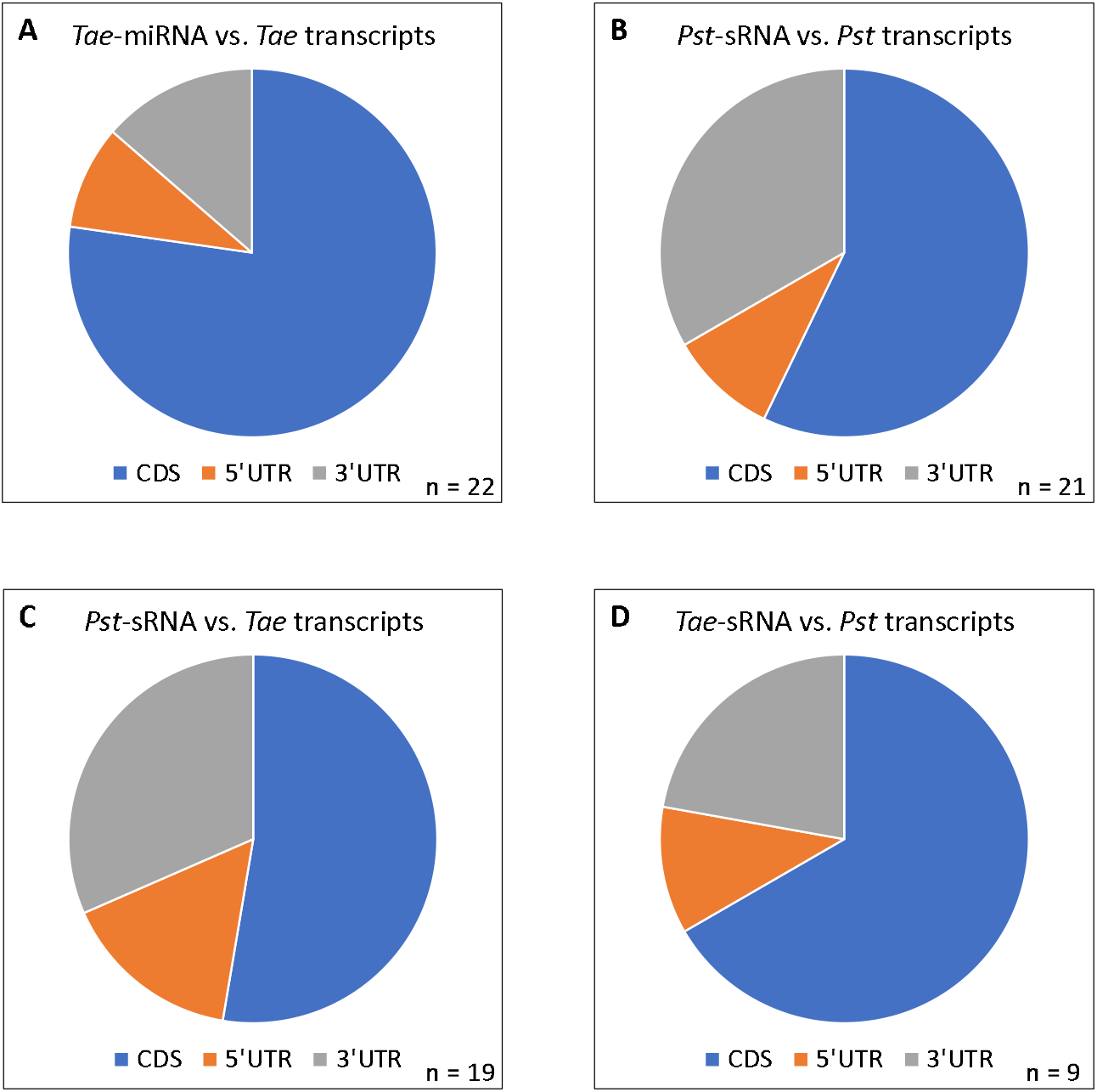
Frequency of observed sRNA-mediated cleavage in the coding sequence (blue), 5’UTR (orange), or 3’UTR (gray) of target transcripts. (**a**) Wheat miRNA vs. wheat transcripts; (**b**) *Pst*-sRNA vs. *Pst* transcripts; (**c**) *Pst*-sRNA vs. wheat transcripts; (**d**) Wheat sRNA vs. *Pst* transcripts.

#### 3.4.3. Fungal sRNAs target fungal transcripts

PARE identified 20 *Pst*-sRNAs with 21 total *Pst* target transcripts. All *Pst*-sRNAs were 20-22 nt in length. Like the overall sRNA library, the majority (70%) began with 5’ uracil (**Table 5**). The genomic origin of most *Pst*-sRNAs overlapped protein-coding genes, which were also often inverted repeat regions (**Supplementary File S9**).

**Table 5.**
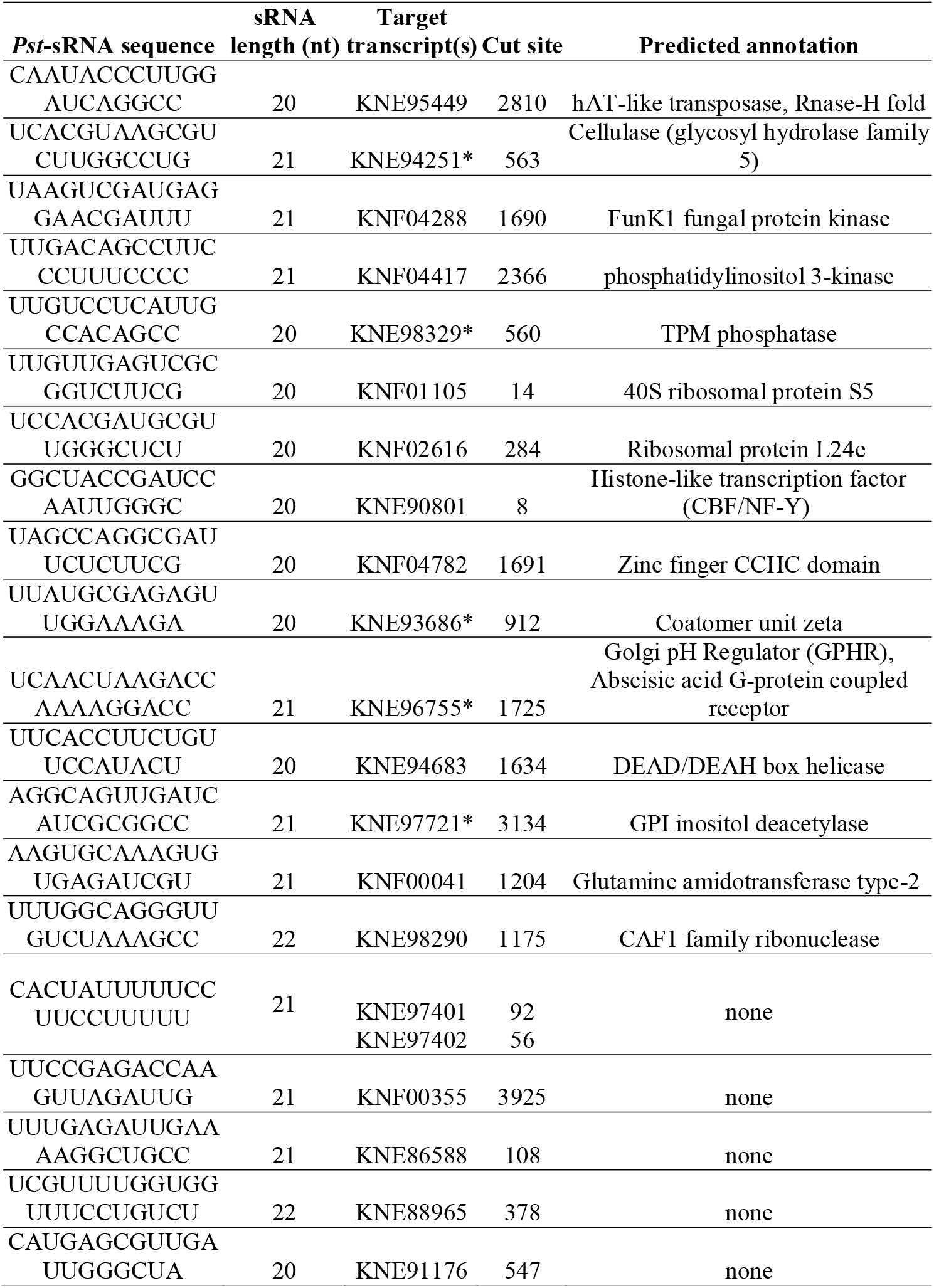
PARE: fungal sRNA vs. fungal transcripts. *Predicted protein contains a signal peptide or transmembrane topology.

Most target transcripts (57%) had cut sites within the coding region, 10% were in the 5’UTR, and 33% were in the 3’UTR (**Figure 6B**). Independent RNA-seq data indicated that 18 out of 21 target genes identified by PARE are expressed in wheat seedlings at seven days post infection (**Supplementary File S8**, [21]). Protein BLAST searches of fungal transcripts targeted by *Pst*-sRNAs found that all but two have a close homolog in another *P. striiformis* isolate, P-130 (**Supplementary File S7**). Furthermore, 81% of target genes have a homolog in at least one of four other cereal rust species: *P. graminis, P. triticina, P. coronata, and P. sorghi*, indicating that most are conserved within the genus. HMMER searches of translated target sequences revealed that they are predicted to encode proteins involved in protein translation, signal transduction, and pathogenesis (**Table 5**). A 21 nt *Pst*-sRNA targets transcript KNF04288, encoding a 703 AA protein homologous to FunK1 protein kinases from many fungal species. This finding suggests that protein phosphorylation is a target for fungal gene regulation and supports previous bioinformatic predictions that *Pst*-sRNA targets are enriched for kinases [18]. Another kinase on the list, Phosphatidylinositol 3-kinase, was found to be important for development and virulence in the plant pathogenic fugus *Fusarium graminearum* [53]. A transposase gene from the hAT transposon superfamily (**Figure 5C**) supported previous findings that transposable elements can be both the source and the target of fungal small RNAs [54]. Ribosomal proteins appear in **Table 5**; small RNAs are known to regulate translation in animals by both upregulating and downregulating ribosomal biogenesis [55, 56]. Since these results suggest that this may also be true in fungi, further molecular genetic research (perhaps using a gene homolog in a model fungus that is easier to transform) could shed light on how ribosomal protein expression is regulated. Five proteins carry a signal peptide and/or transmembrane domain, including KNE94251. This transcript encodes a 445 AA protein with a glycoside hydrolase family 5 catalytic domain found in cellulose-degrading enzymes (**Figure 5D**). The protein may be secreted as one of a cocktail of cellulolytic effectors to help degrade the plant cell wall. Fungal milRNAs are known to target fungal virulence factors [57]. It can benefit pathogens to silence effectors when they might be recognized and elicit effector-triggered immunity in the host. For example, pathogen-derived small RNAs silence an avirulence factor in the Oomycete *Phytophthora sojae*, enabling infection on plants which carry the corresponding resistance gene [58].

#### 3.4.4. Fungal small RNAs target wheat transcripts

PARE data showed evidence of cross-kingdom gene silencing, i.e., that pathogen-derived small RNAs degrade wheat transcripts. Seventeen *Pst*-sRNA sequences target 19 wheat transcripts (**Table 6**). Long inverted repeat regions of the fungal genome near predicted genes were the most common source of *Pst*-sRNAs with wheat targets (**Supplementary File S9**). Most target transcripts (68%) showed independent evidence of being expressed in wheat seedlings during stripe rust infection (**Supplementary File S8**, [21]. Predicted annotations for the polypeptides encoded by these target transcripts show several that are likely to be involved in plant defense against pathogens (**Table 6**, **Supplementary File S7**). A 21 nt *Pst*-sRNA specifically degrades the transcript produced by a gene located on wheat Chromosome 2D (**Figure 5E**). The transcript codes for a 547 amino acid protein with an N-terminal nucleotide binding domain and C-terminal Leucine-rich repeats, two hallmarks of plant resistance genes. NLR proteins recognize pathogen effectors either directly or indirectly to activate effector-triggered immunity [59]. Another target was a transcript coding for glutathione S-transferase, an enzyme family that is important in managing oxidative stress during both biotic and abiotic challenge and is often induced upon microbial infection [60]. Several transcription factors that are often associated with the induction of defense-related genes also appear on this list, including two with a basic leucine zipper (bZIP) domain, homeobox domain, and lateral organ boundaries domain [61] (**Figure 5F**).

**Table 6.**
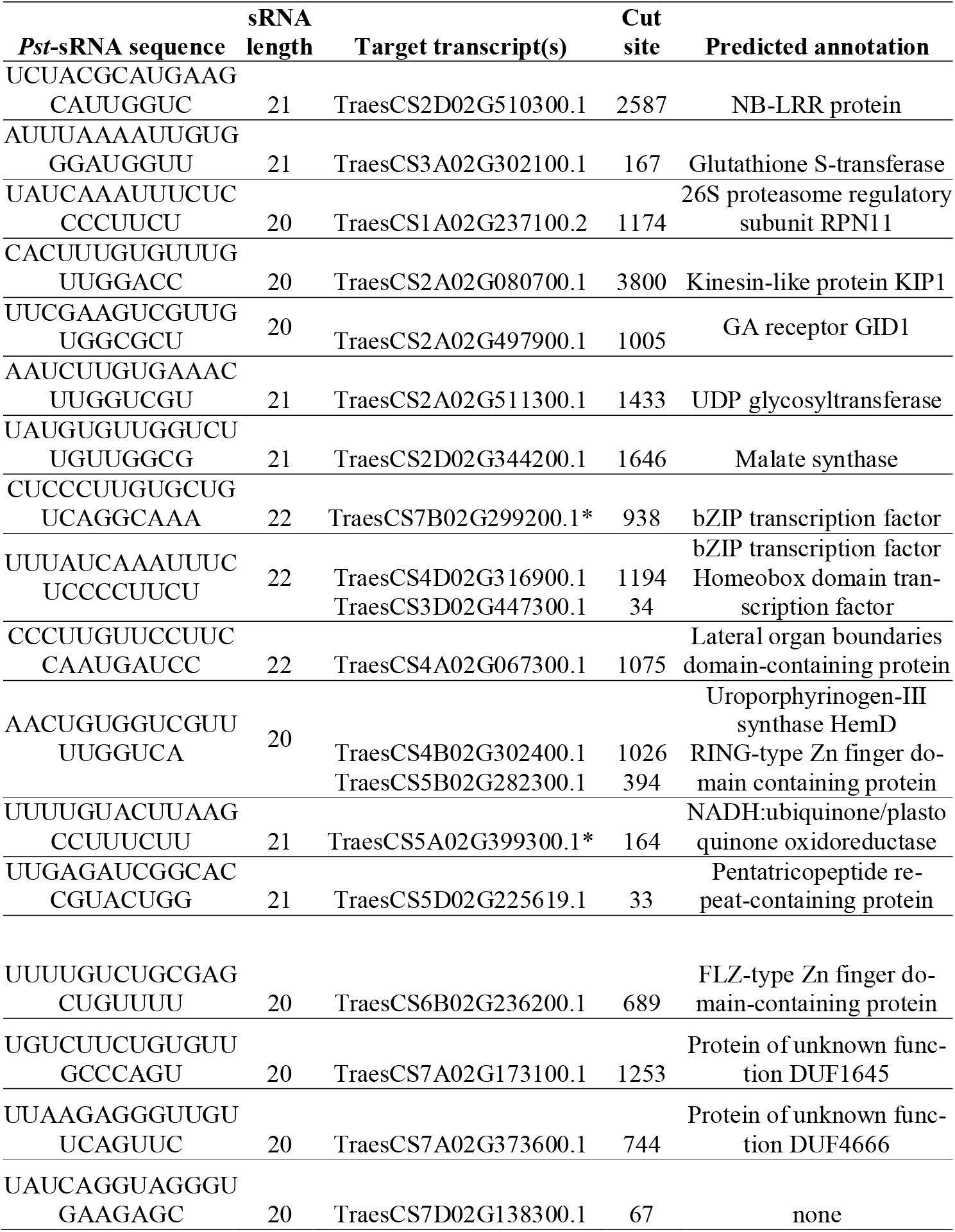
PARE: *Pst-*sRNA vs. wheat transcripts. *Predicted protein contains a signal peptide or transmembrane topology.

Wang and colleagues (2017) reported that the microRNA-like sequence *Pst*-milR1 cleaves the wheat *SM638* transcript, which encodes a β-1,3-glucanase belonging to the Pathogenesis-Related 2 (PR2) gene family [47]. However, we were unable to corroborate this finding. PARE sequencing tags did map to the putative target transcript (TraesCS3D02G475200.1), but there was no evidence of site-specific cleavage (**Supplementary File S10**). No other sequences in the wheat transcriptome showed evidence of being cleaved by *Pst*-milR1 either.

Ten out of 19 cut sites were found within the coding regions of target wheat transcripts, while nine were in the 5’ or 3’ untranslated regions (**Figure 6C**). The proportion of target sites in the UTRs has interesting evolutionary implications. A crucial plant defense transcript targeted by a pathogen sRNA would be expected to experience diversifying selection, with a tendency to remove the complementary sRNA binding site and avoid being degraded. Presumably, this would be more costly if the cut site is in the coding region, since any mutations would be subject to the stipulation that protein function be maintained. UTR sequences, however, are thought to be less functionally constrained. Therefore, over evolutionary time, one might expect cross-kingdom sRNA-target pairs to be biased towards coding sequences. The location of within-species target sites varies widely among species. In many animals including humans, nearly all miRNA target sites are in the 3’UTR, while in *Drosophila* they are split nearly evenly between coding sequence and 3’UTR [62]. In plants, they are typically in the coding regions, with a minority in the 3’UTR [63].

Fungal small RNAs functioning as effective virulence factors are expected to silence the expression of target host transcripts during infection [15]. On the other hand, some plant defense-related genes are induced during infection and may experience a compensatory upward pressure on expression [64]. Thus, the rate of net transcript accumulation associated with cross-kingdom gene silencing may change from repression to induction over time. Target transcripts may temporarily drop in expression and then recover, or simply not be induced as much as they otherwise would be. Reanalyzing a previous RNA-seq time course of wheat seedlings infected with stripe rust [21] provided preliminary support for this idea. Eight out of 19 target transcripts (42%) showed a distinctive expression pattern during early infection: increasing from 0 to 1 dpi; dropping sharply from 2-5 dpi; and finally increasing again from 5-7 dpi (**Figure 7**). The remaining transcripts either did not show this pattern, or had insufficient expression (< 1.0 RPKM) to observe a clear trend. Although more work is needed to assess the effects of *Pst*-sRNAs on host susceptibility, these results suggest that many cross-kingdom gene targets identified by PARE result in perturbation of target transcripts *in planta*.

**Figure 7.**
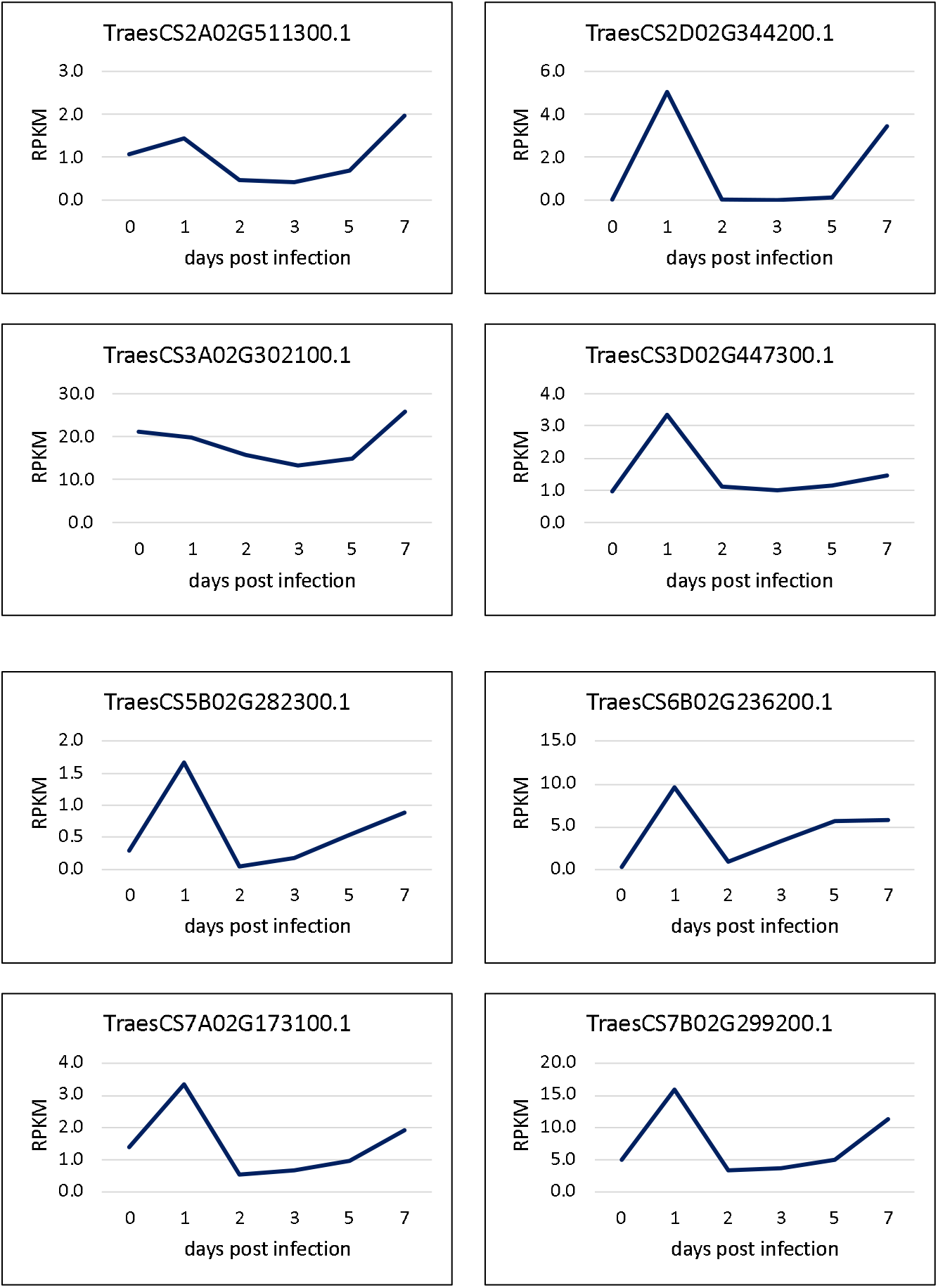
Expression pattern of select wheat transcripts targeted by *Pst*-sRNA over time. Values are mean reads per kilobase per million (n = 3). Raw data from [21].

#### 3.4.5. Wheat small RNAs target fungal transcripts

Cross-kingdom gene silencing was also observed in the opposite direction: plant-derived small RNAs targeting fungal genes. If translocated into the haustoria of *P. striiformis*, these sequences could interfere with various cellular processes by silencing complementary pathogen transcripts. Eight candidate *Tae*-sRNAs targeted a total of nine *Pst* transcripts (**Table 7)**. One previously described wheat microRNA [65], tae-miR10518, was among the nine. Six of nine transcripts had cut sites within the coding sequence; three were cleaved in the UTRs (**Figure 6D**). All but one target gene in *P. striiformis* isolate PST-78 also had a close homolog in the related isolate PST-130 sequences; eight out of nine had significant homology in at least one of four other *Puccinia* spp.: *P. graminis, P. triticina, P. coronata, and P. sorghi* (**Supplementary File S7**). All target genes except one (KNE87418) were expressed in an RNA-seq dataset of stripe rust infecting wheat seedlings (**Supplementary File S8**, [21]). A 22 nt wheat siRNA targets a pair of transcripts from a glycosyl hydrolase gene (PSTG_04871, transcripts KNF02052 and KNF02053). Previous work indicated that glycosyl hydrolase family 26 members are essential for pathogenicity when partial silencing using a fragment of PSTG_04871 expressed by VIGS reduced disease development in wheat seedlings [66]. Similar experiments with this gene, or a *P. graminis* homolog (PGTG01215), indicated the genes were essential for pathogenicity of the stripe rust, stem rust (*P. graminis*) and leaf rust (*P. triticina*) fungi. The *P. graminis* transcript is highly expressed in haustorial cells and protein products are predicted to be secreted, possibly to make sugars more available from mannans, a common carbohydrate target of family 26 glycosyl hydrolases. Thus, wheat siRNA may target these genes to interfere in the primary function of haustorial cells, nutrient acquisition.

**Table 7.**
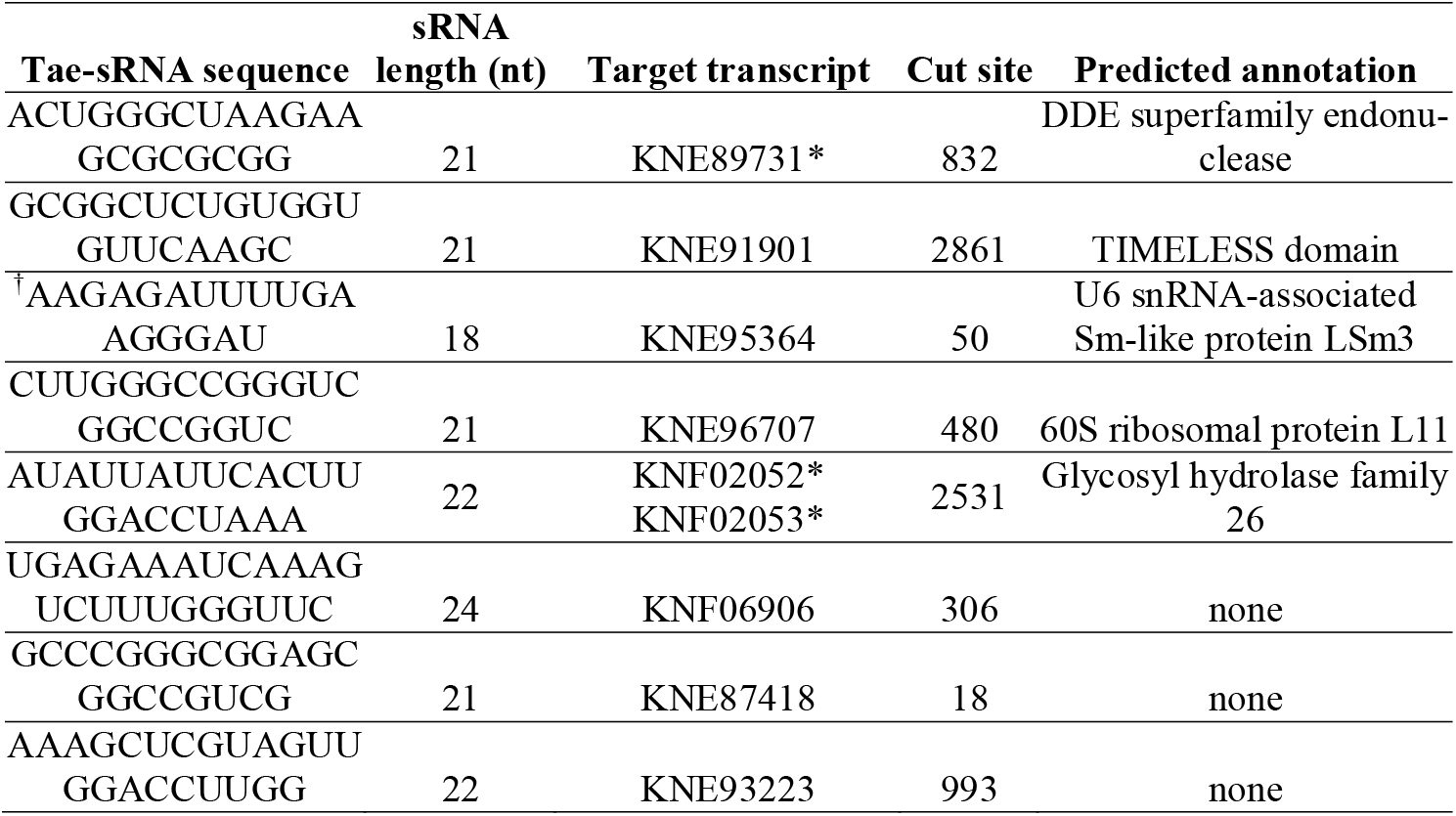
PARE: Wheat sRNA vs. *Pst* transcripts. *Predicted protein contains a signal peptide or transmembrane topology. ^†^Known as microRNA tae-miR10518 [65].

A 21 nt siRNA targets KNE96707, a transcript coding for one protein component of the large ribosomal subunit (**Figure 5G**). As discussed previously regarding *Pst*-sRNA vs. *Pst* transcripts, ribosomal protein expression may be natively regulated by fungal small RNAs. For plants, hijacking this process to inhibit ribosomal biogenesis might be a defense against fungal disease. Ribosomal biogenesis is a target of modern antifungal compounds [67]. Another wheat siRNA targets the transcript of an N-terminal TIMELESS domain-containing protein (**Figure 5H**). Originally discovered as a circadian clock component in *Drosophila melanogaster*, these proteins associate with the DNA replication fork, and are required for the stress response in yeast [68]. It may benefit the plant defense response to disrupt the expression of such fungal proteins via post-transcriptional gene silencing [69].

## 4. Conclusions

This study examined both sides of an inter-kingdom conversation between a staple crop plant and one of its most pervasive pathogens. Among wheat sequences, we observed 24 nt hc-siRNA and 35 nt tRNA and rRNA-derived small RNAs that are differentially expressed upon infection with *P. striiformis*. These are potentially important in the response to stress from fungal pathogens. PARE results in wheat recapitulated known miRNA targets and identified plant small RNAs that appear to direct the slicing of *P. striiformis* transcripts. The annotations of fungal target genes included those involved in protein biosynthesis as well as a previously identified glycosyl hydrolase effector protein. However, this study was based on a compatible interaction with a single susceptible host genotype. Querying the expression of these sRNAs in a broader range of host genotypes could determine how much variation exists in their expression, and whether they are associated with different levels of resistance. If so, successful natural examples of host-induced gene silencing may be used as genetic markers for resistance or even as templates for pathogen control via RNAi-based fungicides [8].

In *P. striiformis*, we described origins and targets of abundant 19-23 nt small RNAs. Several loci met the criteria for MIRNA annotation. No strong homology to *Pst-MIRNA* loci were found in other *Puccinia* spp., suggesting that they are not conserved across diverse fungi, but may have originated since the speciation of *P. striiformis*. An additional locus for the previously-described *Pst-*milR1 revealed that it derives from a long inverted repeat region near predicted genes, displaying a phased pattern of 21 nt reads; additional examples of phasiRNA generation were documented. RT-PCR or Northern analysis using probes specific to these regions could distinguish whether the RNA precursors result from run-on transcription beyond the border of one gene, or rather from natural antisense transcription on both DNA strands. Additional research is needed to find out whether they are analogous to trans-acting siRNAs (tasiRNAs) in plants: secondary siRNAs triggered by miRNA-guided cutting of a primary *TAS* transcript. Small RNAs from long inverted repeat regions have targets in *P. striiformis*, indicating likely regulation of genes involved in transposons, kinases, and a plant cell wall degrading enzyme. The fact that the latter gene is apparently targeted for post-transcriptional silencing poses questions about the temporal and spatial regulation of effectors. Since *P. striiformis* can infect both wheat and *Berberis spp*., it would be interesting to collect RNA from infected barberry plants to compare whether expression levels of pathogenesis-related genes and small RNAs are tailored to each host. One could also study the developmental effects of *Pst*-sRNAs by comparing PARE libraries made from spores and germ tubes versus libraries from infected plant tissue.

One assumption of this study is that post-transcriptional gene silencing in *P. striiformis* is mechanically similar to plants, namely that repression occurs via mRNA slicing between the tenth and eleventh nucleotide of a guide sRNA loaded into an Argonaute protein. Although structural and functional studies of fungal Argonautes from yeast and *Magnaporthe oryzae* give every indication that this is true [69, 70], it is possible we have overlooked transcript cleavage at other positions, or have ignored translational silencing modes that are not detectable with the present method.

Cross-kingdom targets of *Pst-*sRNAs among wheat transcripts included homologs of well-known plant defense genes: an NLR, glutathione-S-transferase, and various biotic stress-related transcription factors. These targets are consistent with the tendency of biotrophic pathogens to suppress the induction of MAMP- and effector-triggered immunity. However, a limitation of this study is that PARE can only identify sRNA-target pairs; it cannot by itself show whether the interaction results in a meaningful decrease in protein levels. Gene expression data indicated that many, but not all cross-kingdom target transcripts experience a dip in expression during early infection, suggesting that induction and suppression occur in a complex back-and-forth dynamic. More transcriptomic and/or proteomic work is necessary to confirm the expression of sRNAs and their targets during infection. To further investigate this intriguing result, a future time course should include mock-inoculated control plants at each time point, as well as quantification of the involved small RNAs over time to observe reciprocal expression of *Pst*-sRNAs and their targets. Showing functional significance is further complicated by the huge diversity of resistance genes such as NLRs in the wheat genome [72]. Also, the present approach cannot show conclusively that small RNAs or their precursors are translocated from pathogen cell to host cell and vice versa, for example via extracellular vesicles [73].

If a fungal small RNA increases virulence by silencing an important plant defense gene, then transforming that plant with an mRNA carrying a target mimic or slicing-resistant target site may lead to an increase in resistance. These sites could be adjusted to suit any pathosystem and updated frequently to circumvent one aspect of pathogen evolution. CRISPR base-editing techniques in cereals have advanced to the degree that transcripts with synonymous mutations that are resistant to slicing have been developed [73, 74]. Further work will determine if durable gains in resistance might be engineered in this way.

## Supplementary Materials

The following supporting information can be downloaded at: www.mdpi.com/xxx/s1, Figure S1: RT-PCR to confirm sample quality; Table S2: oligos used; Table S3: most abundant 35 nt wheat reads and their annotations; Table S4: *Pst*-MIRNA loci; Figure S5: nucleotide and protein sequence alignment of PSTG_08424 and PSTG_08425; Table S6: phased *Pst*-sRNA loci in inverted repeats; Table S7: PARE sRNA-target pair analysis; Table S8: Expression data for PARE target genes; Table S9: Genomic loci producing *Pst*-sRNAs that cleave *Pst* and wheat transcripts; Figure S10: Raw PARE output for the putative target of *Pst*-milR1.

## Author Contributions

NM: methodology, investigation, software, visualization, validation, data curation, writing-original draft; NM and SH: conceptualization, funding, data analysis, writing–review and editing; SH: project administration, supervision. All authors have read and agreed to the published version of the manuscript.

## Data Availability Statement

Sequencing reads for small RNA and PARE libraries are deposited at the NCBI Sequence Read Archive, BioProject PRJNA289147, BioSamples SAMN10968590 through SAMN10968597.

## Acknowledgments

We thank Weiwei Du, Ben Liu, Derek Pouchnik, and Mark Wildung of the WSU Molecular Biology and Genomics Core Facility for sequencing assistance and advice. We thank Kamiak High-Performance Computing staff for bioinformatics resources. Rui Xia and colleagues at South China Agricultural University assisted with sRNA locus annotation.

