## Supplementary File 1 for "Small RNAs target native and cross-kingdom transcripts on both sides of the wheat stripe rust interaction"

**Figure S1. RT-PCR to confirm sample quality.** cDNA from Uninfected wheat (U1-U4) and Infected wheat (I1-I4) amplified using primers specific to wheat Glyceraldehyde-3-phosphate dehydrogenase (GAPDH) and fungal-specific *Pst*-Actin transcripts.

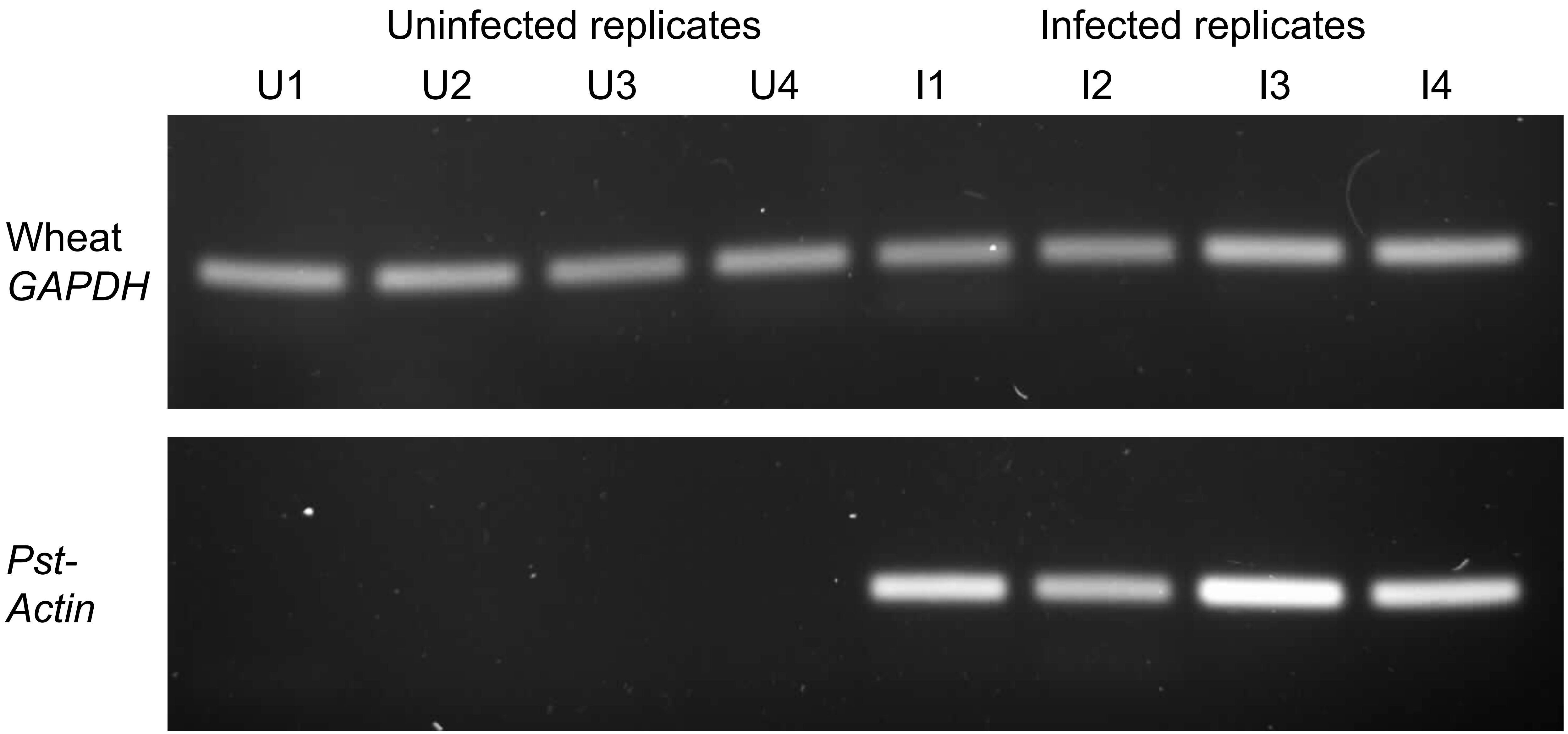
